# Shared and idiosyncratic coding regimes coexist in macaque IT

**DOI:** 10.64898/2026.08.25.746485

**Authors:** Qi Lu, Xinyi Xiong, Yang Li, Hongfei Jiang, Pinglei Bao, Shiming Tang

## Abstract

Population-level measurements portray object representations in primate inferotemporal cortex (IT) as smooth, low-dimensional, and predictable by deep neural networks (DNNs). However, it remains unclear whether this structured population-level picture is representative of the full diversity of its constituent neurons. Here, we used Neuropixels 2.0 probes to record large populations of well-isolated units from an fMRI-localized face patch in macaque anterior IT while monkeys viewed more than 3,000 natural images. Single-unit responses were substantially sparser and more heterogeneous than multi-unit activity (MUA). Sparse neurons collectively formed a higher-dimensional code, supported efficient image identification, and responded later than broadly tuned neurons. Partly independently of sparseness, some neurons exhibited reliable feature randomness: their stimulus preferences were reproducible across repeated presentations but discontinuous across DNN feature spaces, rather than reflecting trial-to-trial response variability or noise. These neurons were virtually uncorrelated with the surrounding population despite being located within the same face patch. By contrast, MUA responses were denser, more correlated, lower-dimensional, and better predicted by DNNs. Together, these findings suggest a dual coding architecture in anterior IT: shared low-dimensional structure supports category generalization, whereas sparse and reliably feature-random single-neuron responses expand the representational space and support efficient identification of individual visual inputs. Response sparseness and reliable feature randomness may therefore constitute fundamental computational resources for object recognition.

## Introduction

Population-level measurements have revealed a remarkably structured view of object representation in the primate inferotemporal cortex (IT) (Bao et al., 2020; DiCarlo & Cox, 2007; Kriegeskorte et al., 2008). Neural responses exhibit smooth, category-related geometry (Bracci & Op de Beeck, 2016; Cadieu et al., 2014; Chang & Tsao, 2017; DiCarlo et al., 2012), can often be described by a relatively small number of dimensions (Chang & Tsao, 2017; Lehky et al., 2014; Lehky & Tanaka, 2016), and are increasingly well predicted by deep neural networks optimized for visual recognition (Khaligh-Razavi & Kriegeskorte, 2014; Schrimpf et al., 2020; Yamins et al., 2014). Together, these findings portray IT as a structured representational space governed by relatively systematic relationships among visual inputs.

Whether this population-level description adequately captures the response properties of individual neurons, however, remains unclear (Hung et al., 2005; Majaj et al., 2015; Rust & DiCarlo, 2010). A regular population geometry does not necessarily imply that all constituent neurons follow the same organizing principles. Shared response structure may coexist with substantial differences in neuronal selectivity, tuning organization, and correspondence to computational models. Moreover, commonly used measurements such as fMRI and multi-unit activity (MUA) combine signals across multiple neurons (Buzsáki, 2004; Kriegeskorte et al., 2008; Majaj et al., 2015; Papale et al., 2025). Such measurements provide robust estimates of locally shared activity but may be less sensitive to response properties that vary across individual neurons. Consequently, the apparent structure of an IT representation may depend partly on the scale at which neural activity is measured.

Resolving this issue requires characterizing both the diversity of single-neuron responses and the population structure that emerges from them. This has been challenging because conventional single-unit studies typically sample limited numbers of neurons or stimuli, whereas large-scale population measurements often lack single-neuron resolution. Comparisons across different recording techniques are further complicated by differences in animals, recording locations, and stimulus sets. A direct test therefore requires large-scale single-unit recordings, broad sampling of visual inputs, and comparison of single-unit and pooled signals derived from the same underlying neural activity.

Here, we used Neuropixels 2.0 probes (Steinmetz et al., 2021) to record large populations of well-isolated units from an fMRI-localized face patch in macaque anterior IT (AIT) while monkeys viewed more than 3,000 natural images. From the same recordings, we extracted single-unit activity (SUA) and MUA and compared their response distributions, correlation structure, population dimensionality, information content, temporal dynamics, and predictability from DNN features. We also examined whether reliable neuronal preferences were organized continuously within the feature spaces learned by current DNNs.

Single-unit responses revealed substantially greater sparseness and heterogeneity than MUA. Sparse neurons collectively formed a higher-dimensional code, supported efficient image identification, and responded later than broadly tuned neurons. In addition, some neurons exhibited reliable stimulus preferences that were discontinuous across DNN feature spaces, partly independently of their response sparseness. These neurons contributed little to the locally shared response structure and were poorly captured by current DNNs. In contrast, MUA emphasized a denser, more correlated, lower-dimensional, and more model-predictable component of the neural response. These findings show that the structured representation observed at the population level is genuine but does not capture the full diversity of the underlying single-neuron code.

## Results

### Large-scale single-unit recordings from macaque anterior IT during natural-image viewing

We first established a large-scale single-unit dataset from fMRI-localized face-selective regions in macaque anterior inferotemporal cortex. In two monkeys, anterior face-selective cortex was identified with fMRI and used to guide the placement of a transparent recording window and Neuropixels 2.0 probe insertions (Fig. 1A, Fig. S1A,B). A total of one or two four-shank Neuropixels probes were inserted, either singly or in parallel, through the perforated window into the targeted AIT region (Fig. S1C,D), allowing simultaneous sampling of single-unit activity across cortical depth and nearby cortical locations. Tissue drift was limited during recording, and post hoc drift correction enabled stable isolation of large populations of single units (Fig. S2A,B). Subsequent analysis of neuronal selectivity for faces confirmed the fMRI localization of the face patch (Fig. S1E,F, see methods) (Bao et al., 2020; Freiwald & Tsao, 2010; Shi et al., 2026; Tsao et al., 2006).

**Fig. 1.**
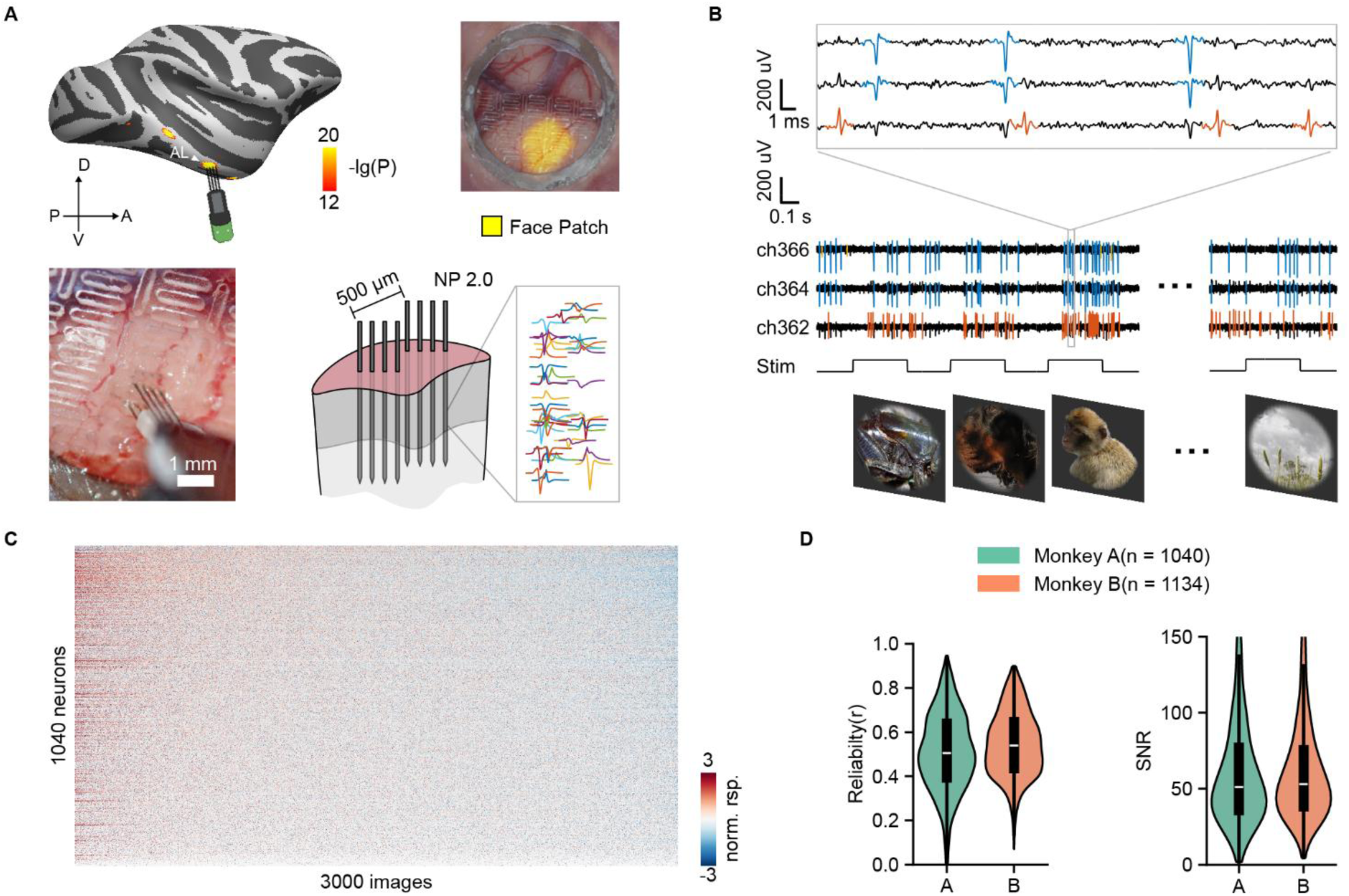
Large-scale recording of macaque AIT neuronal responses to natural images. **A,** fMRI-guided Neuropixels recordings from an anterior inferotemporal (AIT) face patch. Upper left, face-selective activation projected onto the NMT v2.0 macaque cortical template. Upper right, alignment of the fMRI-defined face patch with the cortical surface visible through the recording chamber using vascular landmarks. Lower left, photograph of two implanted Neuropixels 2.0 probes. Lower right, schematic of probe implantation and example spike waveforms across recording sites. **B,** Visual stimulation and electrophysiological recording. Top, example extracellular voltage traces; blue and orange waveforms indicate spikes assigned to two different single units. Middle, multichannel spiking activity during stimulus presentation. Bottom, example natural images from ImageNet. **C,** Response matrix for 1,040 single units recorded from Monkey A across 3,000 natural images. Neurons are ordered by increasing sparseness and images by decreasing mean population response. Responses were z-scored across images separately for each neuron. **D,** Distributions of response reliability (left) and spike-waveform signal-to-noise ratio (SNR; right) for single units from Monkey A (*n* = 1,040) and Monkey B (*n* = 1,134). Reliability was quantified as the split-half correlation across repeated stimulus presentations and corrected using the Spearman–Brown formula. Black boxes indicate the median and interquartile range.

During recordings, monkeys passively viewed thousands of natural images while maintaining fixation. Images were drawn from ImageNet (Deng et al., 2009) and related natural-image collections, spanning multiple visual categories including animals, faces, objects, and natural scenes (Fig. 1B). To sample both the broad natural-image space and stimuli that effectively drove AIT neurons, the final stimulus set consisted of approximately two-thirds randomly selected natural images and one-third response-optimized images selected from an initial screening procedure (Fig. S3; Table S1, see Methods). Each image was repeated 6–8 times, allowing us to estimate the reliability of image-evoked responses (Fig. S3).

Spike sorting of the Neuropixels recordings using Kilosort4 (Pachitariu et al., 2024), followed by additional quality control (Fabre et al., 2023), yielded large populations of visually responsive single units in both animals (see Methods). Example raw traces showed robust stimulus-evoked spiking across channels (Fig. 1B). For each unit, image-evoked responses were quantified as baseline-corrected firing rates, computed by subtracting the mean activity during the −100 to 0 ms pre-stimulus baseline from the mean activity during the 100–300 ms post-stimulus response window. This response window was chosen based on the average AIT response time course (Fig. S2E). The resulting image-by-neuron response matrix revealed substantial diversity in visual responses across AIT neurons (Fig. 1C), with some neurons responding broadly to many images and others responding selectively to a restricted subset.

The final dataset contained 1,040 visually responsive single units from Monkey A and 1,134 from Monkey B. Across both animals, units showed reliable responses across repeated image presentations and robust signal-to-noise ratios (Fig. 1D). The median refractory period violation rate was 0.80% (Fig. S2C), indicating excellent spike sorting quality and high-quality SUs. The median missed-spike ratio was 0.18% (Fig. S2D), indicating reliable spike waveforms. This dataset therefore provided a high-quality basis for directly comparing the representational structure revealed by single-unit activity with that inferred from locally pooled multi-unit signals.

### Single-unit responses reveal sparse image selectivity hidden by multi-unit signals

We next asked whether well-isolated single units and locally pooled multi-unit signals reveal different response structures in macaque AIT. Neuropixels 2.0 provides dense recording sites, allowing us to isolate single-unit activity through spike sorting while also extracting MUA from the same raw recordings. We therefore compared three neural signal types: quality-controlled SUA from Neuropixels 2.0 recordings, MUA extracted from the same Neuropixels raw signals using a standard MUA processing pipeline (Majaj et al., 2015), and a previously published Utah-array MUA dataset (Papale et al., 2025) recorded during natural-image viewing (Fig. 2A).

**Fig. 2.**
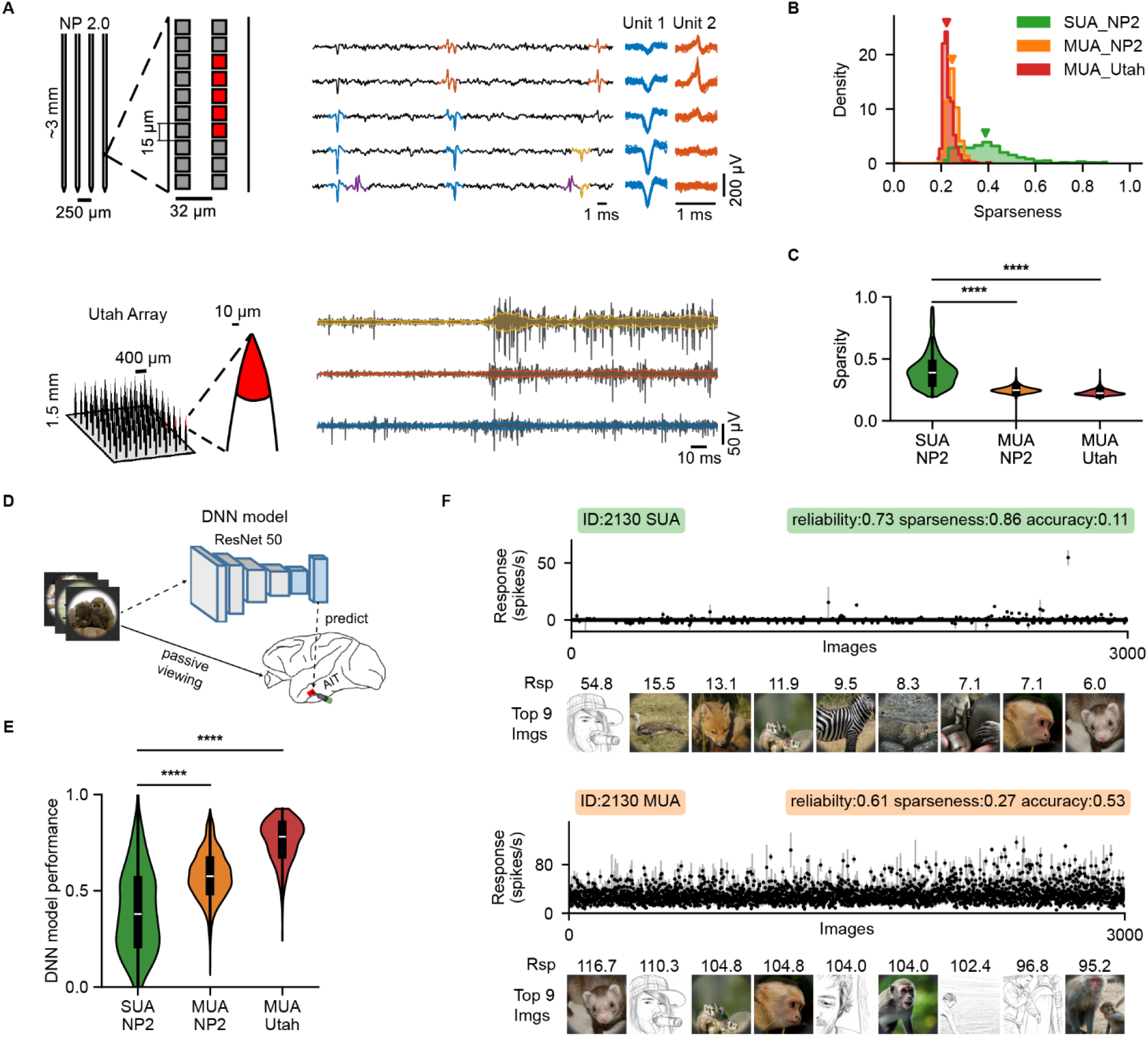
SUA responses span a broader range of sparseness and are less predictable by DNNs than MUA responses. **A,** Electrode configurations and example signals from Neuropixels 2.0 (NP2; top) and Utah array recordings (bottom). Left, electrode schematics and enlarged views illustrating recording-site spacing. Middle, example multichannel extracellular voltage traces; colored spike waveforms in the NP2 traces were assigned to different units by spike sorting, whereas the colored curves superimposed on the Utah array traces show the signal envelopes used to compute multi-unit activity (MUA). Upper right, multichannel spike waveforms of two example NP2 single units. **B,** Distributions of response sparseness for NP2 SUA, NP2 MUA, and Utah array MUA. Arrowheads indicate medians. NP2 SUA comprised units isolated by spike sorting and subsequent quality control; NP2 MUA was extracted from the corresponding raw NP2 signals using the MUA processing pipeline; and Utah MUA was obtained from a previously published dataset of Utah array responses to natural images. **C,** Comparison of response sparseness across NP2 SUA, NP2 MUA, and Utah MUA. **D,** DNN encoding-model pipeline. ResNet-50 features extracted from the presented images were used to predict neural responses through a two-layer multilayer perceptron, using separate training, validation, and test image sets. **E,** Comparison of noise-ceiling-normalized DNN prediction performance across NP2 SUA, NP2 MUA, and Utah MUA. **F,** Responses of an example sparse SUA (top) and the MUA signal recorded at the corresponding site (bottom) across 3,000 images. Points and error bars indicate the mean ± s.e.m. response across repeated presentations of each image. The nine most effective images for each signal are shown below, with the corresponding mean firing rates indicated in spikes s⁻¹. Image examples are schematic illustrations generated by GPT-5.6 Sol to represent the content of the stimuli. Human face images have been replaced with cartoon illustrations. Labels above each plot report response reliability, sparseness, and DNN prediction performance. In **C** and **E**, black boxes indicate the median and interquartile range. Between-group comparisons were performed using two-sided Mann– Whitney *U*-tests. \**P* < 0.05, \*\**P* < 0.01, \*\*\**P* < 0.001, and \*\*\*\**P* < 0.0001.

To quantify response selectivity across the natural-image set, we computed Hoyer sparsity (Hoyer, 2004) for each unit (see Methods). Higher Hoyer sparsity indicates that a unit responds strongly to a smaller fraction of images and therefore provides a statistical measure of image selectivity in our stimulus set. This comparison revealed a clear difference in response sparseness.

SUA exhibited a broad sparseness distribution, extending into a high-sparseness range rarely observed in either Neuropixels-derived or Utah-array MUA, and was significantly sparser than both MUA signals (Fig. 2B,C). To assess the generality of this pattern, we extended the comparison to the independent Neuropixels-based Triple-N dataset (Y. Li et al., 2026) and a second Utah-array MUA dataset (Majaj et al., 2015). The same pattern was observed across datasets: Neuropixels SUA showed broader, higher-sparseness distributions, whereas MUA from both Neuropixels and Utah arrays remained narrower and shifted toward lower sparseness (Fig. S4). Thus, the greater sparseness of SUA primarily reflected the distinction between isolated single-neuron responses and locally pooled activity rather than differences between datasets or recording platforms.

Consistent with this interpretation, averaging SUA responses within progressively larger spatial neighborhoods around a selected MUA channel produced responses that increasingly resembled MUA, with the strongest MUA-SUA-average correlation emerging at an intermediate spatial scale of approximately 100– 200 μm (Fig. S5A,B) (Buzsáki, 2004). Increasing the averaging radius also progressively reduced response sparseness, supporting the view that local pooling transforms sparse single-unit responses into denser MUA-like signals (Fig. S5C,D).

We next tested whether these differences in response structure affected model predictability. We built a DNN-based encoding model by extracting image features from a pretrained ResNet-50 (He et al., 2016) and fitting a two-layer perceptron to predict neural responses using train-validation-test cross-validation (Fig. 2D).The best-performing ResNet-50 layer was conv5_block1_add_bn (layer 153; Fig. S6), and prediction performance was quantified as the prediction correlation normalized by each unit’s noise ceiling.

DNN encoding performance was markedly lower for Neuropixels SUA than for either Neuropixels MUA or Utah MUA (Fig. 2E; median normalized performance: Neuropixels SUA, 0.38; Neuropixels MUA, 0.58; Utah MUA, 0.78). Moreover, DNN predictability increased as SUA responses were averaged over progressively larger spatial neighborhoods (Fig. S5F), indicating that local pooling makes AIT responses more aligned with DNN feature representations. Example responses showed the same pattern: a sparse SUA responded strongly to only a few images and was poorly predicted by the model, whereas the MUA signal from the same location showed dense responses and substantially higher predictability (Fig. 2F, SUA: sparseness 0.86, accuracy 0.11; MUA: sparseness 0.27, accuracy 0.53). Thus, DNN models better captured pooled MUA signals than isolated single-neuron responses, suggesting that SUA exhibits a highly selective tuning structure that is partially obscured by MUA. Together, these results show that local pooling obscures sparse single-neuron selectivity in macaque AIT. While MUA signals reveal a dense, model-predictable component of AIT activity, SUA shows a broader and more selective range of image tuning that is substantially less aligned with current DNN-based encoding models.

### Sparse single-unit responses form a richer and higher-dimensional population code

The greater sparseness of SUA responses does not necessarily imply a simpler population code. If different single neurons respond sparsely to different subsets of images, the resulting population activity may span a richer, higher-dimensional representational space. For example, some SUA showed highly selective tuning for specific visual features, including particular facial configurations, face viewpoints, or individual object identities. (Fig. S7). We therefore next asked whether the sparse response structure revealed by SUA was associated with differences in population organization relative to MUA.

We first visualized population response matrices by randomly sampling 100 units from each signal type and plotting their responses to the 100 images that elicited the strongest average responses across the sampled population (Fig. 3A–C). Consistent with the sparseness analysis, SUA responses appeared more heterogeneous across neurons, with individual units showing more selective and less overlapping response patterns. In contrast, both Neuropixels-derived MUA and Utah-array MUA showed denser and more homogeneous response structure across images.

**Fig. 3.**
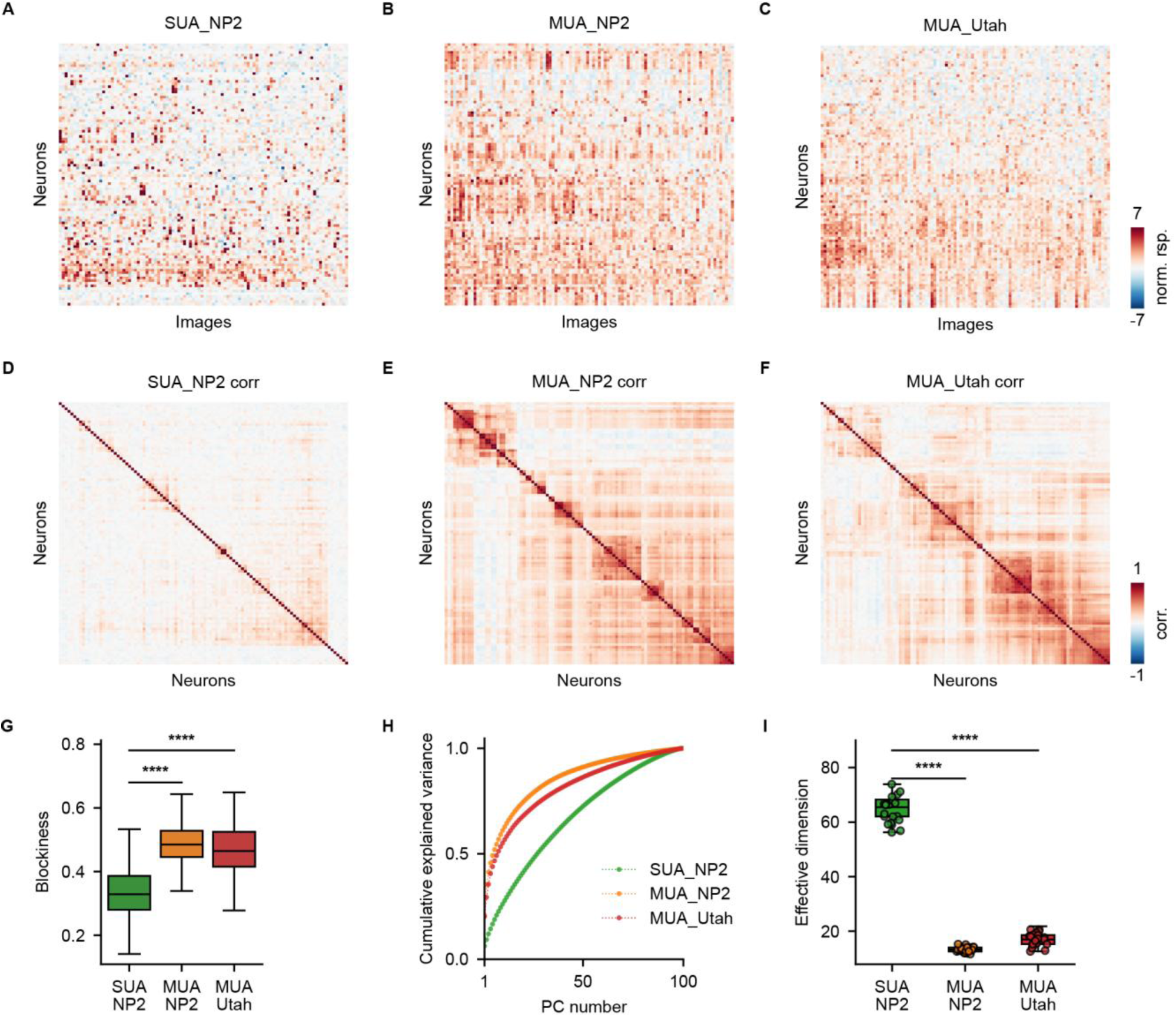
SUA responses exhibit weaker block structure and higher effective dimensionality than MUA responses. **A–C,** Response matrices for 100 randomly sampled response profiles from NP2 SUA (**A**), NP2 MUA (**B**), and Utah array MUA (**C**). For visualization, the 100 images eliciting the highest population-mean responses in each dataset are shown. Response profiles were ordered by response-based clustering, and responses were z-scored across images separately for each profile. **D–F,** Pairwise Pearson correlation matrices for the same response profiles and ordering as in **A–C**, computed using responses to all images. **G,** Comparison of correlation-matrix blockiness across the three datasets. Blockiness was defined as the mean within-cluster correlation minus the mean between-cluster correlation, with cluster membership determined from response-based clustering. **H,** Cumulative variance explained by successive principal components for the response profiles shown in **A–C**, calculated using responses to all images. **I,** Comparison of effective dimensionality across NP2 SUA, NP2 MUA, and Utah array MUA. Effective dimensionality was calculated from 100 randomly sampled response profiles, with sampling repeated 40 times for each dataset. Between-group comparisons were performed using two-sided Mann– Whitney *U*-tests. \**P* < 0.05, \*\**P* < 0.01, \*\*\**P* < 0.001, and \*\*\*\**P* < 0.0001.

To characterize the shared tuning structure within each population, we computed pairwise correlations between response profiles across images. Groups of units with similar image preferences would appear as blocks of elevated correlation in the resulting matrices. Such block structure was prominent in both MUA datasets but substantially weaker in SUA, which also showed lower overall pairwise response correlations (Fig. 3D–F). By contrast, both MUA datasets showed stronger, more clustered correlation structure, indicating that local pooling increased similarity across recorded channels. We quantified this property using a blockiness measure and found that the SUA correlation matrix was significantly less block-like than either Neuropixels-derived MUA or Utah-array MUA (Fig. 3G). Thus, MUA signals appear to compress neural diversity into a smaller number of shared response patterns.

We next asked how these differences in correlation structure affect representational dimensionality. Principal component analysis revealed that SUA required substantially more principal components to explain the same fraction of response variance, whereas MUA responses were captured by a smaller number of dominant components (Fig. 3H). Consistent with this result, the effective dimension (Del Giudice, 2021; Pirkl et al., 2012) of SUA populations was markedly higher than that of both MUA datasets when computed from randomly sampled populations of 100 neurons and repeated across resamples (Fig. 3I). Further linking this difference to local pooling, progressively averaging SUA responses over larger spatial neighborhoods led to a corresponding reduction in effective dimensionality (Fig. S5E). Thus, despite being sparser at the level of individual neurons, SUA populations formed a higher-dimensional code.

Together, these results show that sparse single-neuron responses in AIT do not reflect a low-dimensional or simplified representation. Instead, SUA responses are more heterogeneous, less correlated, and higher-dimensional than MUA signals. Local pooling, therefore, not only reduces apparent sparseness but also compresses the representational richness of the underlying single-neuron population.

### Response sparsity and encoding randomness jointly limit DNN model predictability

We next asked why SUA responses were less predictable by DNN encoding models. One contributing factor was response sparseness. Across SUA, model performance decreased with lifetime sparseness (Fig. 4A; *r* = −0.57), indicating that neurons responding strongly to fewer images were generally more difficult to predict. For such units, prediction depends heavily on whether the model can capture the small number of images that evoke strong responses.

**Fig. 4.**
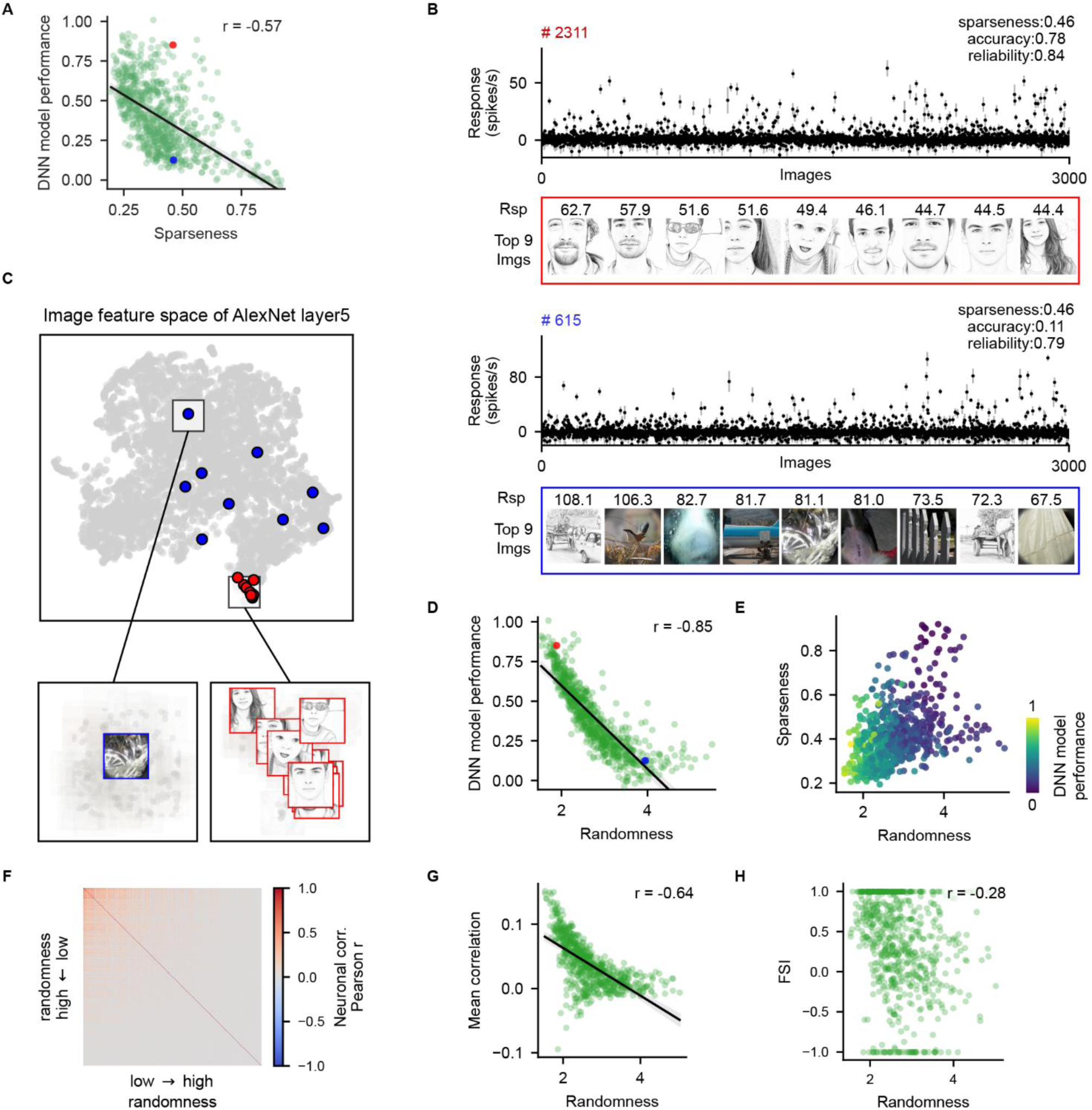
SUA sparseness and feature randomness jointly account for variation in DNN prediction performance. **A,** Relationship between DNN prediction performance and response sparseness across single units. The solid line shows the linear regression fit, and the shaded band indicates its 95% confidence interval (CI). Red and blue points indicate two example units with comparable sparseness but markedly different DNN prediction performance. **B,** Response profiles of the two example units highlighted in **A** across 3,000 images. Points and error bars indicate the mean ± s.e.m. response across repeated presentations of each image. The upper unit showed consistent preferences for faces, whereas the lower unit preferred a visually diverse set of images. The nine most effective images and their corresponding mean responses are shown below each response profile. Labels report response sparseness, DNN prediction performance, and response reliability. **C,** Two-dimensional embedding of the image feature space derived from AlexNet layer 5. Gray points represent all images, whereas red and blue points indicate the preferred images of the low- and high-randomness example units in **B**, respectively. Boxed regions are enlarged below together with the corresponding images. **D,** Relationship between DNN prediction performance and feature randomness. Randomness quantifies the discontinuity of neuronal responses across neighboring images in DNN feature space and does not reflect trial-to-trial response variability. The red and blue points indicate the same example units as in **A–C**. The solid line shows the linear regression fit, and the shaded band indicates its 95% CI. **E,** Joint distribution of response sparseness, feature randomness, and DNN prediction performance. Each point represents one unit, and color indicates noise-ceiling-normalized DNN prediction performance. **F,** Pairwise response-correlation matrix, with units ordered by increasing feature randomness along both axes. Each entry represents the Pearson correlation between the response profiles of two units across all images. **G,** Relationship between feature randomness and the mean response correlation of each unit with all other units, excluding self-correlations. The solid line shows the linear regression fit, and the shaded band indicates its 95% CI. **H,** Relationship between feature randomness and the face-selectivity index (FSI). Reported r values are Pearson correlation coefficients. For image examples in **B** and **C**, human face images have been replaced with cartoon illustrations.

Sparseness alone, however, did not fully explain model performance. Two example neurons had nearly identical sparseness but markedly different prediction accuracies (Fig. 4A, red and blue points). Both neurons exhibited highly reliable response profiles, yet the better-predicted neuron responded preferentially to a coherent set of face images, whereas the poorly predicted neuron preferred a heterogeneous collection of images (Fig. 4B). Thus, DNN predictability appeared to depend not only on how selectively a neuron responded, but also on how its preferred images were organized within the model feature space.

To quantify this organization, we measured the spatial autocorrelation of each neuron’s responses in the AlexNet layer 5 feature space (Krizhevsky et al., 2017; Moran, 1950); (Fig. S8, see Methods). This measure captures the continuity of the mapping from image features to neuronal responses: low feature randomness indicates that nearby images in feature space evoke similar responses, whereas high feature randomness indicates that reliable response preferences are distributed discontinuously across the feature space. Consistent with this distinction, the preferred images of the better-predicted example neuron occupied a compact region of the feature space, whereas those of the poorly predicted neuron were widely dispersed (Fig. 4C). Additional examples confirmed that the measure agreed with the apparent organization of neuronal preferences (Fig. S9). Randomness estimates were strongly correlated across DNN layers (Fig. S10A,B) but only weakly related to neuronal reliability and sparseness (Fig. S10C,D), indicating that feature randomness did not simply reflect response noise or sparse firing. We also computed and compared the randomness in feature spaces from different DNNs, including AlexNet, VGG19 (Simonyan & Zisserman, 2015), ResNet-50, and ViT-B/16 (Dosovitskiy et al., 2020), and found that the randomness based on deep-layer feature spaces was relatively consistent across networks (Fig. S11).

Across the SUA population, feature randomness was strongly negatively correlated with DNN model performance (Fig. 4D; *r* = −0.85), and this relationship was consistent across feature spaces derived from different DNN layers (Fig. S10E). Plotting model performance in the joint space defined by sparseness and feature randomness showed that the best-predicted neurons were concentrated in the lower-left region, where responses were both less sparse and less feature-discontinuous (Fig. 4E). Prediction performance declined as either property increased. A multiple linear regression incorporating both factors closely reproduced the observed variation in DNN predictability (Fig. S12; correlation between fitted and observed performance, *r* = 0.87). Multiple regression revealed distinct contributions of the two factors. Feature randomness remained strongly associated with lower DNN predictability after controlling for sparseness (partial *r* = −0.80, *P* <10^−165^; unique Δ*R*² = 0.433, 95% CI = 0.383–0.480), whereas sparseness made a smaller but significant independent contribution (partial *r* = −0.40, *P* < 10^−28^; unique Δ*R²* = 0.045, 95% CI = 0.027–0.067). Together, they explained 76.1% of the variance in DNN predictability (R² = 0.761), with feature randomness contributing substantially more unique variance. These results suggest complementary constraints on model predictability: sparseness limits the amount of informative response variation available for model fitting, whereas feature randomness limits generalization across nearby images in the model-defined feature space.

Feature randomness was also related to how individual neurons participated in the surrounding population code. When neurons were ordered by randomness, response correlations were strongest among low-randomness neurons and progressively weakened toward neurons with higher randomness (Fig. 4F). Accordingly, a neuron’s mean response correlation with the rest of the recorded population decreased markedly with randomness (Fig. 4G; *r* = −0.64). Highly random neurons were therefore nearly uncorrelated with the shared population response. Randomness was also negatively related to the face selectivity index (Fig. 4H; *r* = −0.28), indicating that neurons with more feature-discontinuous preferences tended to show weaker face selectivity despite being recorded within a face patch.

SUA and MUA also differed substantially in feature randomness (Fig. S13A,B). Linking individual SUA responses with the MUA recorded at their corresponding sites revealed a marked contraction in the sparseness–randomness space: the broadly distributed SUA responses were compressed into the lower-left region occupied by denser and less random MUA responses (Fig. S13C). This shift is consistent with the view that local pooling attenuating sparse, neuron-specific response components while preserving feature-continuous structure shared across neurons. We also visualized the spatial arrangement of neurons and found that neurons with high feature randomness were intermingled with the overall neuronal population (Fig. S14).

Together, these results identify response sparseness and reliable feature randomness as separable properties that jointly constrain the ability of DNNs to predict SUA responses. Feature-random neurons contribute little to the shared, category-selective response structure of the local population, and this component becomes less apparent in pooled MUA measurements.

### Sparse and non-sparse neurons support complementary visual computations and differ in response timing

The preceding analyses identified response sparseness and feature randomness as partly separable properties of single-neuron responses. We next asked whether these properties were associated with differences in the information represented by neuronal populations. We first divided SUA into high- and low-randomness groups and repeatedly sampled 40 neurons from each group. High-randomness populations showed substantially greater effective dimensionality than low-randomness populations (Fig. 5A; Mann– Whitney *U* test, *P* < 0.0001). Thus, although high-randomness neurons were weakly correlated with the surrounding population, their heterogeneous response profiles collectively spanned a richer representational space.

**Fig. 5.**
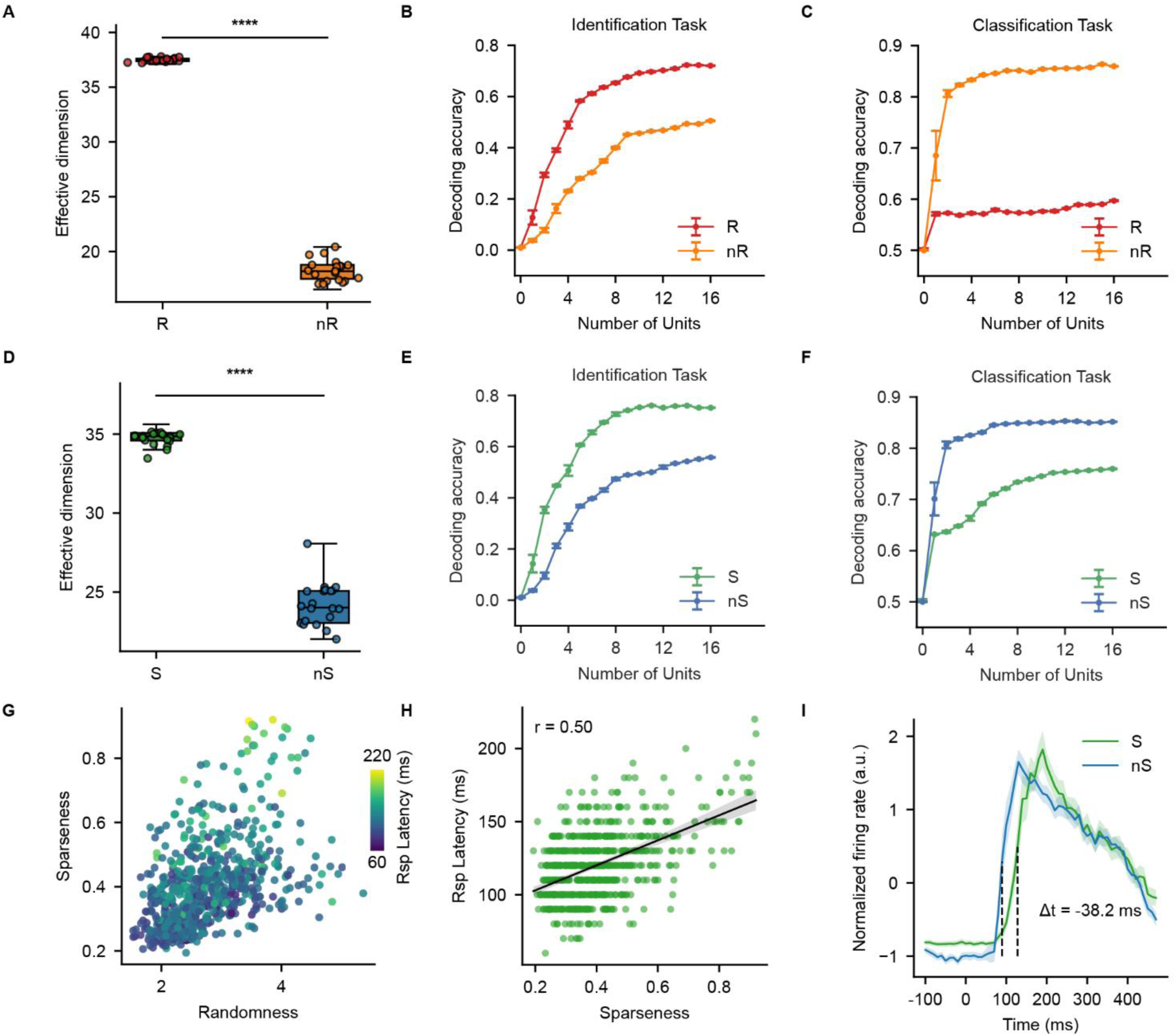
Functional differentiation and temporal differences of single-neuron subsets divided by randomness and sparsity. **A,** Effective dimensionality of populations formed by randomly sampling 40 random (R) or 40 non-random (nR) neurons. **B, C,** Identification (**B**) and classification (**C**) decoding accuracy as a function of the number of random or non-random neurons. **D,** Effective dimensionality of populations formed by randomly sampling 40 sparse (S) or 40 non-sparse (nS) neurons. **E, F,** Identification (**E**) and classification (**F**) decoding accuracy as a function of the number of sparse or non-sparse neurons. **G,** Distribution of response latency in the randomness– sparseness space. Each point represents one neuron, and color indicates its response latency. **H,** Relationship between response latency and sparseness (*r* = 0.50). The solid line shows the linear regression fit, and the shaded band indicates the 95% confidence interval. **I,** Mean stimulus-evoked response time courses of sparse and non-sparse neurons, expressed as z-score-normalized firing rates. Vertical dashed lines mark the group response latencies; sparse neurons responded on average 38.2 ms later than non-sparse neurons. Shaded regions indicate 95% confidence intervals. Comparisons in **A** and **D** used two-sided Mann–Whitney *U*-tests. Asterisks indicate statistical significance: \**P* < 0.05, \*\**P* < 0.01, \*\*\**P* < 0.001 and \*\*\*\**P* < 0.0001.

We next compared the two groups in image-identification and category-classification tasks. High-randomness neurons supported more accurate image identification and reached a given level of performance with fewer neurons than low-randomness neurons (Fig. 5B). The opposite pattern was observed for category classification: low-randomness neurons substantially outperformed high-randomness neurons, particularly at small population sizes (Fig. 5C). Thus, feature-discontinuous responses were not simply unstructured or uninformative. Rather, they efficiently distinguished individual visual inputs, whereas feature-continuous neurons better captured category-level structure shared across images.

Response sparseness was associated with a parallel differentiation. Populations of sparse neurons exhibited higher effective dimensionality than equally sized populations of non-sparse neurons (Fig. 5D; Mann– Whitney *U* test, *P* < 0.0001). Sparse neurons also supported more efficient image identification (Fig. 5E), whereas non-sparse neurons performed better in category classification, including face versus non-face discrimination (Fig. 5F). Therefore, although sparseness and feature randomness describe partly independent aspects of neuronal responses, both were associated with a shift from shared category-level structure toward a higher-dimensional representation of individual images.

We finally examined whether these response properties were related to neuronal response timing. Visualizing response latency across the joint randomness–sparseness space suggested that the clearest temporal variation occurred along the sparseness dimension (Fig. 5G). Across neurons, response latency increased with sparseness (Fig. 5H; *r* = 0.50), indicating that neurons with more selective response distributions generally responded later. This difference was also apparent in the population time courses: the average visually evoked response of sparse neurons was delayed by approximately 38 ms relative to that of non-sparse neurons (Fig. 5I), consistent with previous reports that coarse categorical information emerges earlier than fine-grained identity information in IT responses (Sugase et al., 1999; Suzuki et al., 2006). Thus, the sparse, high-dimensional component that efficiently supported image identification was associated with a later phase of the visual response, whereas broader responses carrying category-level information emerged earlier.

Together, these results reveal a clear functional and temporal dissociation within AIT. Sparse neurons form a higher-dimensional population code, support efficient stimulus identification, and respond later. Non-sparse neurons, by contrast, are lower-dimensional, better support category-level classification, and respond earlier. These findings suggest that AIT contains complementary coding regimes: an earlier, broader code suited for coarse categorization, and a later, sparse code suited for fine-grained visual discrimination.

### Random sparse coding supports a high-dimensional population manifold and sparse identification readout

To examine how random sparse coding shapes population geometry and downstream readout, we compared the measured SUA responses with responses reconstructed using an axis model (Fig. 6A; Fig. S15A). The axis model constrains each neuron’s response to vary smoothly along a single direction in feature space, thereby providing a benchmark for smooth, low-dimensional axis coding. The model captured only a limited portion of the measured response structure across all three signal types, with a median prediction performance of 0.29 for SUA (Fig. S15B–D). At the single-neuron level, the axis-model-fitted responses were concentrated within a narrow region of low randomness and low sparseness, whereas the measured SUA occupied a substantially broader region of the joint randomness–sparseness space (Fig. 6A, top; Fig. 6B). At the population level, measured SUA also exhibited markedly higher effective dimensionality than the axis-model-fitted responses (Fig. 6A, middle; Fig. 6C), indicating that their heterogeneous tuning expanded the population response manifold beyond that captured by smooth axis coding.

**Fig. 6.**
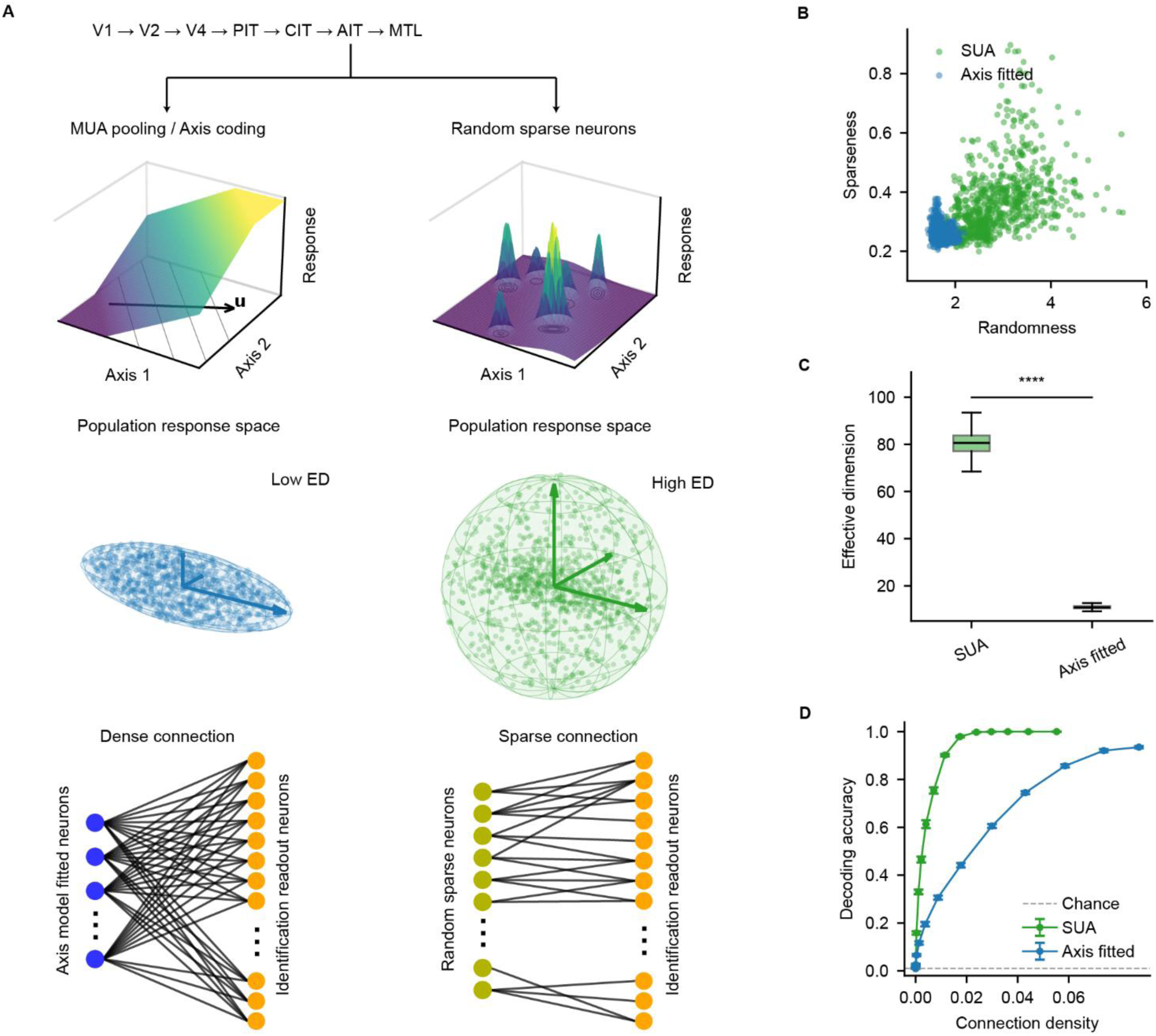
Random sparse coding supports high-dimensional population representations and sparse downstream readout. **A,** Schematic comparison of low-dimensional smooth coding (MUA pooling or axis coding) and random sparse coding at three levels. Top, an axis-model-fitted neuron exhibits smooth tuning along a feature axis, whereas a random sparse neuron responds at multiple isolated locations in feature space. Middle, axis coding produces a low-dimensional population response manifold, whereas random sparse coding produces a higher-dimensional manifold. Bottom, identification from axis-model-fitted responses requires relatively dense readout connections, whereas random sparse responses support identification with sparser connections. **B,** Joint distribution of feature randomness and sparseness for measured single-unit activity and responses fitted by the axis model. Each point represents one neuron. **C,** Effective dimensionality of measured SUA and axis-model-fitted population responses. **D,** Identification decoding accuracy as a function of downstream connection density for measured SUA and axis-model-fitted responses. The horizontal dashed line indicates chance performance. The comparison in **C** used a two-sided Mann–Whitney *U*-test. Asterisks indicate statistical significance: \**P* < 0.05, \*\**P* < 0.01, \*\*\**P* < 0.001 and \*\*\*\**P* < 0.0001.

We next asked whether this expanded population geometry allowed individual images to be identified using fewer downstream connections. We trained identification decoders while systematically restricting the proportion of upstream neurons available to each readout unit (Fig. 6A, bottom). Decoding from measured SUA reached 90% accuracy at a connection density of only 0.01, whereas decoding from axis-model-fitted responses required a density of approximately 0.07 to achieve comparable performance (Fig. 6D). Thus, within the decoder tested here, the high-dimensional population manifold associated with random sparse coding enabled accurate image identification with approximately sevenfold sparser connectivity than the smooth, low-dimensional axis-coded representation.

### Sparse and feature-random units in DNNs preferentially support image identification

We next asked whether response sparseness and feature randomness were unique to biological neurons or were also present in DNN units. We randomly sampled 10,000 artificial units from the conv5_block1_out layer (layer 154) of ResNet-50 and measured their responses to the same natural images presented to the monkeys (Fig. S16A). To exclude largely inactive units, we retained only units whose maximum activation exceeded 10, yielding approximately 5,000 artificial neurons for subsequent analyses. The same measures of feature randomness and lifetime sparseness were then applied to the artificial and biological responses.

Artificial and biological neurons showed substantially overlapping distributions of feature randomness, although the biological population exhibited greater heterogeneity and a more extended high-randomness tail (Fig. S16B). Their sparseness distributions were also centered at similar values, but biological neurons again spanned a broader range and included more extremely sparse responses (Fig. S16C). Thus, sparse and feature-discontinuous response profiles were not unique to AIT neurons, although their diversity was greater in the biological population. We also examined the data sensitivity of the sparseness and randomness calculations using simulated neurons (Fig. S17) and found that 1,600 natural image stimuli were sufficient to accurately estimate both randomness and sparseness.

We next tested whether feature randomness was associated with similar information-processing differences in artificial units. High-randomness artificial units supported more efficient image identification than low-randomness units, reaching high decoding accuracy with fewer units (Fig. S16D). In contrast, low-randomness units consistently performed better in category classification (Fig. S16E). We further evaluated the contribution of different randomness levels to the DNN’s own classification performance by retaining units from successive randomness ranges. Classification accuracy remained relatively stable across the lower four randomness ranges but decreased sharply when only the most-random units were retained (Fig. S16F). Therefore, although high-randomness units carried information sufficient to distinguish individual images, they contributed relatively little to the feature-continuous structure supporting the network’s category classification.

A similar functional differentiation was observed for response sparseness. Sparse artificial units supported substantially more efficient image identification than non-sparse units (Fig. S16G), whereas non-sparse units performed better in category classification (Fig. S16H). Classification based on units from different sparseness ranges was highest for units with intermediate sparseness and declined markedly for the two most-sparse ranges (Fig. S16I). Extremely sparse artificial units therefore retained strong image-specific information but were less effective in supporting the DNN’s category decisions.

Together, these results show that DNNs contain artificial units spanning both feature randomness and response sparseness, and that these properties are associated with an identification-classification trade-off resembling that observed in biological neurons. High-randomness and sparse units efficiently distinguish individual inputs, whereas less-random and moderately sparse units better support category-level classification.

## Discussion

The present study shows that the representational structure inferred from macaque anterior inferotemporal cortex depends strongly on recording resolution. Large-scale Neuropixels recordings revealed that single-unit activity was substantially sparser, more heterogeneous, higher-dimensional, and less predictable from current DNN features than multi-unit activity derived from the same recordings. Progressively averaging SUA across larger spatial neighborhoods reproduced these MUA-like properties. Within the SUA population, response sparseness and reliable feature randomness emerged as partly independent dimensions of neuronal variation. Both were associated with higher population dimensionality and more efficient image identification, whereas broader and more feature-continuous responses better supported category classification. These findings suggest that AIT contains both a shared, category-related component that is emphasized by local pooling and a heterogeneous, image-specific component that is most apparent at single-neuron resolution.

Local pooling does not provide a neutral summary of the underlying neuronal population. Response components shared across nearby neurons add constructively, whereas neuron-specific components are attenuated, providing a parsimonious account of why MUA responses were denser, more correlated, lower-dimensional, less feature-random, and better predicted by DNNs. This does not imply that the structured geometry observed in pooled signals is artifactual. Rather, pooled measurements accurately capture the locally shared component of AIT activity while providing a less complete view of single-neuron variation. This observation clarifies how the smooth, category-related geometry emphasized by population-level measurements can coexist with the highly selective responses documented in individual neurons. Shared structure becomes dominant after local pooling, whereas single-neuron recordings reveal an additional heterogeneous component that expands the representational space. Accurately characterizing population-average geometry therefore does not necessarily imply that the response properties of its constituent neurons have been fully explained.

Response sparseness and feature randomness captured distinct aspects of this single-neuron heterogeneity. Sparseness describes how strongly a neuron concentrates its responses on a small subset of images. Feature randomness instead describes whether its reliable response preferences vary continuously within a model-defined feature space. Neurons with comparable sparseness could differ markedly in feature randomness and DNN predictability, and both properties explained unique variance in model performance. Moreover, randomness was only weakly related to response reliability, indicating that it did not simply reflect trial-to-trial noise. We therefore use the term reliable feature randomness to describe reproducible but feature-discontinuous tuning.

This randomness is necessarily relative to the feature spaces used to measure it. A neuron that appears discontinuous in AlexNet or ResNet feature space may be organized with respect to visual variables that those models do not represent. High randomness should therefore not be interpreted as an absence of structure in an absolute sense. Instead, it identifies a component of neuronal tuning that is not locally smooth within current model representations. The finding that high-randomness neurons were nearly uncorrelated with the surrounding population despite being recorded within a face patch, further suggests that they lie outside the dominant category-related organization of the local population.

Despite their weak correspondence to this shared structure, sparse and high-randomness neurons were not uninformative. Populations enriched for either property exhibited higher effective dimensionality and supported image identification with fewer neurons. Low-randomness and non-sparse populations, in contrast, better supported category classification. Thus, the same response properties that make individual neurons difficult to predict from shared feature spaces can increase the diversity and image discriminability of the population code. AIT may therefore balance two complementary computational regimes: a correlated and feature-continuous representation that supports generalization across category members, and a more heterogeneous representation that separates individual visual inputs.

These regimes should not be interpreted as two discrete neuronal classes. Sparseness and randomness varied continuously across the SUA population and were only partly correlated with one another. Some neurons were sparse but feature-continuous, whereas others exhibited feature-discontinuous preferences without extreme sparseness. Both properties can expand population dimensionality, but they do so in different ways: sparseness reduces the number of stimuli driving each neuron, whereas feature randomness reduces the redundancy among neurons by distributing their preferences differently across feature space.

The extreme image selectivity of some sparse neurons is reminiscent of grandmother-cell accounts (Gross, 2002; Quiroga, 2012). However, selectivity across the images used here does not establish a one-neuron–one-object code. Demonstrating such a code would require showing invariant responses across transformations, viewpoints, contexts, and exemplars of the same identity. Our results demonstrate extreme image selectivity and efficient population-level identification but cannot determine whether the preferred images reflect invariant object identities or particular visual configurations. We therefore interpret these neurons as the sparse end of a distributed code while leaving open the possibility that some exhibit more invariant tuning.

The temporal results provide an additional clue to how this heterogeneous code develops. Response latency increased continuously with sparseness, and sparse neurons responded approximately 38 ms later on average than non-sparse neurons. This pattern is consistent with an early representation dominated by broadly shared visual structure, followed by a later stage that increases stimulus specificity. Local inhibition could contribute to such a transformation by suppressing common response components and allowing only particularly strong or distinctive inputs to remain above threshold. Recurrent excitation, feedback, heterogeneous feedforward convergence, and intrinsic neuronal dynamics provide alternative explanations. Moreover, the presence of sparse and feature-random units in feedforward DNNs shows that these response properties do not, by themselves, require recurrent or inhibitory mechanisms. Establishing their biological origin will require causal perturbations and circuit-level measurements.

One possible origin of feature randomness is heterogeneous and partly idiosyncratic connectivity. Different combinations of feedforward inputs could generate neuron-specific feature conjunctions that are reproducible within an individual system but not aligned with the dominant representational axes of other brains or independently trained networks. Experience-dependent plasticity and synaptic pruning could subsequently refine and stabilize some of these configurations. This possibility is conceptually related to neuronal group selection, or Neural Darwinism, in which developmental variation generates diverse circuit repertoires and experience selects neuronal ensembles that contribute to behavior (Edelman, 1987, 1993). Because selection operates on functional ensembles rather than enforcing one-to-one correspondence between individual neurons, different systems could achieve similar population-level functions through distinct microscopic solutions. Reliable feature randomness may therefore reflect structured, system-specific variation shaped by connectivity and learning history rather than noise in neuronal responses

The relationship between recording scale and DNN predictability also qualifies how model–brain correspondence should be interpreted. DNN performance increased as SUA responses were progressively averaged, indicating that current models preferentially capture response components shared across nearby neurons. Strong prediction of MUA or fMRI therefore does not necessarily imply accurate prediction of the constituent single neurons. Evaluating biologically adequate models will require testing not only whether they reproduce pooled representational geometry, but also whether they capture the distributions of single-neuron sparseness, feature continuity, correlation, dimensionality, and temporal dynamics.

At the same time, the DNN analyses argue against a simple biological-versus-artificial distinction. ResNet-50 contained artificial units spanning a broad range of sparseness and feature randomness, and these properties were associated with an identification–classification trade-off resembling that observed in AIT. Sparse and high-randomness artificial units efficiently distinguished individual images, whereas less-random and moderately sparse units better supported category classification. Biological neurons nevertheless exhibited broader and more extreme distributions of these properties. Thus, current DNNs may instantiate the same general coding principles but differ from AIT in how extensively they use them.

The presence of feature-random units in both systems also suggests a different interpretation of poor cross-system predictability. Shared, low-dimensional response components are likely to align across independently trained models and biological brains because they reflect common regularities of the visual environment and classification objective. High-dimensional, unit-specific preferences need not align in the same way: different networks—or a network and the brain—may implement useful stimulus separation along different axes. A biological neuron may therefore be poorly predicted by a particular DNN even when that DNN contains its own reliable feature-discontinuous units. Feature randomness may represent system-specific expansion of the representational space rather than meaningless noise, although decoding performance alone does not establish that these units are causally used for recognition.

Several limitations constrain this interpretation. Recordings were obtained from an fMRI-localized face patch in anterior IT, and it remains unknown whether the same organization extends across other IT regions. Feature randomness was evaluated relative to a finite set of DNN feature spaces and could decrease with models that better capture the relevant neuronal variables. Image-identification decoding demonstrates that information is available in these populations but not that downstream circuits use it. Finally, spatial averaging and MUA provide controlled tests of local pooling but do not reproduce all properties of larger-scale signals such as fMRI. Addressing these questions will require recordings across multiple IT regions, richer tests of invariance, comparisons with more diverse computational models, and causal circuit manipulations.

In summary, the apparent organization of macaque AIT depends strongly on measurement scale. Local pooling emphasizes correlated, category-related, and model-predictable structure while attenuating sparse and reliably feature-discontinuous single-neuron responses. These heterogeneous responses expand population dimensionality and efficiently distinguish individual visual inputs, whereas broader and more feature-continuous responses better support category generalization. The coexistence of these components in both AIT and DNNs suggests that sparseness and reliable feature randomness provide computational resources for balancing generalization with the separation of individual visual inputs.

## Methods

### Animals and surgeries

All experimental procedures were conducted in accordance with the guidelines of the Institutional Animal Care and Use Committee (IACUC) of Peking University Laboratory Animal Center and were approved by the Peking University Animal Care and Use Committee (LSC-TangSM-3). Two adult male rhesus macaques (*Macaca mulatta*; subjects A and B, aged 4–6 years) were used in this study. Prior to experiments, animals were trained to adapt to head fixation and to perform high-precision visual fixation tasks until stable performance was achieved (fixation error < 1°).

All surgeries were performed under general anesthesia in strictly sterile conditions. Postoperative care included analgesia, antibiotic administration, and continuous health monitoring throughout recovery. Each animal underwent three sequential surgeries: implantation of a headpost system, skull preparation for anterior inferotemporal (AIT) access, and implantation of an electrophysiological recording chamber.

The first surgery involves implantation of the headpost system. A midline skin incision was made to expose the skull, and the surgical site was prepared. Burr holes were drilled using a dental drill, and titanium bone screws were implanted as anchoring points for securing MRI-compatible headposts. The headposts were then firmly attached to the skull using dental acrylic cement, providing long-term mechanical stability for head fixation during subsequent behavioral training and fMRI experiments.

Before the second surgery, functional MRI-guided localization of face-selective patch was performed (Fig. S1A, B). According to the localization, a portion of skull and overlying muscle tissue above the target region in AIT was removed to prepare for chamber implantation. In the same surgery we reconstruct the headpost by implanting three titanium headposts on the skull (two on the forehead and one on the back of the head). A T-shaped steel frame was subsequently attached to the posts to stabilize head position during electrophysiological recordings.

In the third surgery, performed at least two weeks later, a 20-mm craniotomy was made over the temporal cortex to target the superior temporal sulcus (STS) region. The skull and dura were opened to expose the cortical surface. The recording chamber design was based on that described by Li et al. (2017), with two modifications: (1) the glass coverslip was fabricated with closely spaced obround slots to accommodate the insertion of four Neuropixels 2.0 shanks, and (2) silicone membranes were incorporated on both sides of the coverslip to provide a sealed barrier and reduce the risk of infection. Specifically, a custom-fabricated glass coverslip (15 mm in diameter, 0.30 mm in thickness) was prepared with obround slots measuring 1.25 mm × 0.25 mm. The outer and inner surfaces of the glass were covered with silicone membranes 50 μm and 20 μm in thickness, respectively. The glass coverslip was secured within a 10-mm-diameter titanium ring using silicone adhesive. A ring-shaped GORE membrane (12 mm inner diameter, 31 mm outer diameter) was then attached to the titanium ring to form a sealed window unit. The window unit was gently positioned over the cortical surface, with the GORE membrane inserted beneath the dura. The titanium ring was secured to the skull using dental acrylic, thereby forming a stable recording chamber (Fig. 1A). The chamber was further reinforced with hardened dental acrylic and covered with a protective metal shield to prevent damage when the animals were returned to their home cages. Additional details of the surgical procedures and chamber design have been described previously (M. Li et al., 2017).

### Behavior paradigm

Animals were head-fixed and performed passive fixation tasks during recordings. The animal was required to maintain fixation within a 1° circular window around a central fixation point (a small white dot measuring 0.1°) to obtain juice rewards. Eye position was monitored with an infrared eye-tracking system (ISCAN) at 120 Hz.

#### Functional MRI paradigm

In fMRI experiments, a block design was employed. Each trial consisted of 500 ms stimulus-on and 500 ms stimulus-off periods, with 24 trials per block. Each functional run contained 8 stimulus blocks and 9 fixation-only baseline blocks, yielding a total duration of 408 s.

Visual stimuli were displayed on an LCD screen (refresh rate 60 Hz, resolution 1024 × 768) at a viewing distance of 100–120 cm. For functional localization, stimuli included faces, bodies, objects, and scrambled imaged presented in separate blocks (24 images per block; stimulus diameter 8°).

#### Electrophysiology behavioral task

The natural image stimuli used in the large dataset were sourced from ImageNet (Deng et al., 2009), specifically ILSVRC2012 and 8 synsets from the person subtree. The original images were cropped, resized and masked to create round patches measuring 330 pixels (10 degrees) in diameter with soft fade-off. The visual stimuli were generated using the ViSaGe system (Cambridge Research Systems Ltd., UK) and displayed on a 17-inch LCD monitor (Acer V173, refresh rate 80 Hz, resolution 1280 × 960) positioned 45 cm away from the animal.

For schematic illustration in this paper, these stimuli were transformed into cartoon-style graphics using GPT-5.6 Sol. All generated illustrations were manually verified by the authors to ensure they accurately represented the key features of the original stimuli. These illustrations were used solely for visual demonstration in this article.

The macaques were trained to maintain fixation on a small fixation dot (diameter 0.1°). Each stimulus was presented for 300 ms following a pre-fixation period of 200 ms. Trials were aborted if fixation deviated beyond 1°. After every five successful trials, animals received juice rewards.

### MRI acquisition and processing

The fMRI experiment was conducted at the Center for MRI Research at Peking University using a 3T Siemens Prisma MRI scanner. Data acquisition was performed with a custom-built 8-channel surface coil. Prior to functional scanning, high-resolution T1-weighted 3D structural images were acquired for each macaque using a 3D-MPRAGE sequence with the following parameters: repetition time (TR) = 2300 ms, echo time (TE) = 3.8 ms, flip angle = 9°, number of slices = 224, field of view (FOV) = 128 mm × 128 mm, voxel size = 0.5 mm × 0.5 mm × 0.5 mm. Whole-brain functional scans were obtained using an echo-planar imaging (EPI) sequence with parameters: TR = 2000 ms, TE = 20 ms, flip angle = 80°, slices = 27, FOV = 96 mm × 96 mm, voxel size = 1.5 mm × 1.5 mm × 2 mm. To improve the signal-to-noise ratio, monocrystalline iron oxide nanoparticles (MION; Molday ION, 0.26 mL/kg, BioPAL) were injected into the monkey’s femoral vein prior to the functional experiment (Bao et al., 2020).

Surface reconstruction based on anatomical volumes was performed using FreeSurfer (Dale et al., 1999) after skull stripping with the Brain Extraction Tool (FSL, University of Oxford). Functional volumes were analyzed using the FreeSurfer Functional Analysis Stream (Reuter & Fischl, 2011). Motion correction and distortion correction were applied to the functional volumes based on the acquired field maps. The resulting data were then analyzed using a standard general linear model. For the face contrast condition, the mean signal across all face stimulus blocks was compared with the mean signal across body stimulus blocks. All contrasts were assessed using unpaired two-tailed t tests, and no correction for multiple comparisons was applied to the *P* values. After obtaining the face activation regions, they were registered to the NMT template (Jung et al., 2021) using 3dNwarpApply, and the results are shown in Figure 1A.

### Electrophysiological recording

We recorded neuronal responses from the targeted area using Neuropixels 2.0. Each probe contains four shanks (250 μm spacing), 384 recording channels, and 5120 selectable low-impedance TiN electrode sites (12 × 12 μm², impedence of 148 ± 8 kilohms at 1 kHz). Sites are distributed along a 10-mm shank (70 × 24 μm cross-section), with two columns of sites spaced 32 μm apart and 15 μm center to-center spacing along the length of the shank (Steinmetz et al., 2021).

A custom probe insertion system was developed to ensure stable electrophysiological recordings. During recording sessions, the macaque was head-fixed. One or two Neuropixels probes were secured onto a probe holder via dovetail mounts. For recordings using two probes, the probes were positioned with a center-to-center spacing of 500 μm (Fig. 1A). The probe holder was attached to a micromanipulator, which was mounted onto the animal’s T-shaped steel frame using custom-fabricated steel components. The micromanipulator was carefully adjusted to align the probe trajectory perpendicular to the obround slots in the glass coverslip of the recording chamber. The probes were then advanced vertically, penetrating the silicone membrane through the obround slots and into the cortex to a target depth of 3–5 mm, thereby covering the cortical depth range of the AIT region.

After insertion, the probe holder was fixed to the chamber base using dental acrylic, and reference grounding was connected to the headframe. A copper mesh shield was added for electrical shielding. Typically, we waited 30–60 min for stabilization of the probe to minimize drift and ensure stable signal.

Following probe stabilization, 384 recording channels were selected from the 5,120 available electrode sites on each Neuropixels 2.0 probe. For simultaneous recordings with two probes, 768 channels were selected from the combined 10,240 electrode sites and streamed through a single headstage.

Electrophysiological signals from Neuropixels 2.0 probes were acquired through a PXI-based data acquisition module (IMEC) integrated into a PXIe chassis (PXIe-1071, National Instruments). Full-band neural signals (0.5 Hz–10 kHz) were digitized at 30 kHz with 14-bit resolution and continuously recorded using SpikeGLX software (Bill Karsh; https://github.com/billkarsh/SpikeGLX).

To synchronize neural recordings with visual stimulation, stimulus onset signals generated by the ViSaGe system were delivered to the digital input channel of the IMEC acquisition system and recorded simultaneously with the neural signals to provide precise temporal alignment between electrophysiological recordings and visual stimulus events. The neural signal processor (Cerebus system, Blackrock Microsystems) recorded stimulus onset events, image identifiers, and eye position signals, allowing us to accurately align electrophysiological signals with individual visual stimuli.

### Spike sorting and neural response analysis

Electrophysiological data were stored by SpikeGLX in binary format. Raw electrophysiological signals were preprocessed using Kilosort4 (https://github.com/MouseLand/Kilosort/tree/main) (Pachitariu et al., 2024), including high-pass filtering, common-mode noise removal, and drift correction. Following preprocessing, spike sorting was performed using Kilosort4 to detect and cluster action potentials into putative single units. The spike detection thresholds were set to 11 for universal templates and 9 for learned templates. These thresholds were set higher than the default values in Kilosort4 to improve the reliability of isolated spike waveforms.

Following automated sorting, units were manually curated using the open-source software Phy2 (https://github.com/cortex-lab/phy). Units were merged or split based on multiple quality criteria, including spike waveform similarity, drift patterns, and cross-correlogram features. Quality of sorted units was assessed automatically using BombCell (Fabre et al., 2023).

To generate peri-stimulus time histograms (PSTHs) for each unit in response to visual stimuli, firing rates were calculated using a 10-ms sliding window. To quantify visual responsiveness, firing rates were calculated separately during the OFF period (−100 to 0 ms before stimulus onset) and the ON period (100–300 ms after stimulus onset). Stimulus-evoked responses were quantified as the increase in mean firing rate from the OFF period to the ON period.

Custom MATLAB scripts and open-source MATLAB tools (Neuropixel Utils) were used for synchronization of electrophysiological data and behavioral event signals, spike timing analysis, computation of unit-level visual responses, and stringent quality control of recorded units.

#### Unit inclusion criteria

Only units satisfying three inclusion criteria were used for subsequent analyses. First, Kilosort4-classified multi-unit activity (MUA) units were excluded. Unit quality was assessed based on contamination rate estimated from refractory period violations (RPVs) relative to Poisson expectation, and units with contamination rates >20% were removed. Second, units were required to exhibit significant visual responsiveness, defined as a response >10 spikes/s to at least one stimulus. Third, units were required to demonstrate reliable responses across repeated presentations. For each unit, the Pearson correlation between each repetition and the across-repetition mean response profile was calculated. Repetitions with correlations <0.5 were discarded, and only units retaining more than half of their original repetitions were included.

### MUA analysis

To enable direct comparison with SUA, we applied a standard MUA extraction pipeline (Majaj et al., 2015) to the raw signals acquired with Neuropixels 2.0 probes. For each recording channel, raw voltage traces were first band-pass filtered between 250 Hz and 7.5 kHz. Multi-unit spikes were then detected by identifying threshold crossings of negative voltage deflections exceeding three times the standard deviation of the raw voltage signal. Spike events were detected independently for each recording channel, and MUA responses were quantified based on the increase in mean firing rate during the 100–300 ms period after stimulus onset relative to baseline.

In Figure S6, we utilized an alternative approach to simulate MUA responses by pooling adjacent single units. Specifically, for each NP2 recording site, simulated MUA responses were generated by averaging the responses of SUA units located within circular regions centered on the corresponding recording site. The spatial pooling radius was varied across 5, 20, 40, 60, 80, 100, 150, 200, 250, 300, 350, 400, 450, and 500 μm.

### Stimuli screening

The visual stimulus protocol consisted of two stages. In the first stage, we identified site-specific preference images through an initial stimulus screening procedure, selecting 1,000 images (20 sessions) or 800 images (8 sessions). These were then combined with 2,000 or 1,600 session-shared randomly selected natural images, respectively, to form the final stimulus sets of 3,000 or 2,400 images for large-scale recording. Each image was presented 6–8 times.

During the initial stimulus screening phase, 6,000–10,000 randomly selected color natural images were sequentially presented to the macaques for one repetition. For each single unit, an image was considered a preference stimulus if it elicited a response exceeding half of the unit’s maximum response and evoked a firing rate of at least 30 Hz. To ensure balanced sampling across units, the maximum number of preference images contributed by each unit was initially limited to 1000/*n*, where *n* denotes the number of recorded units. This contribution limit was progressively increased until the target number of preference images was reached.

In the large-scale recording phase, the selected site-specific preference images were combined with the session-shared randomly selected natural images to form the final stimulus set. Each image was presented 6–8 times, with image order independently randomized across repetitions, while neuronal responses were recorded. Across the experiments, the dataset comprised 28 recording sessions, with responses from thousands of units collected to these natural image stimuli.

### Quantification of single-neuron visual response characteristics

Python v.3.9 and MATLAB R2022a were used for analysis.

#### Reliability

For the analyses presented in Figures 2–5, we included only units with high response reliability. For each unit, responses were randomly divided across repetitions into two independent subsets. The mean response to each stimulus was calculated separately for each subset, and the Pearson correlation between the two resulting stimulus-response profiles was used as the split-half reliability. This procedure was repeated 100 times with different random splits. The correlation coefficients (r) obtained across iterations were averaged, and the resulting value was corrected using the Spearman–Brown correction to obtain the final reliability estimate.

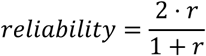

Units with reliability values below 0.6 were excluded from analyses in Figs. 2–5.

#### Face selectivity

We measured face selectivity using the session-shared random natural images. The face selectivity index (FSI) (Tsao et al., 2006) was defined for each unit as:

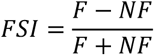

where *F* represents responses to face images and *NF* represents responses to non-face images. The resulting FSI values were clipped to the range of −1 to 1. Because neuronal responses were expressed as baseline-subtracted firing rates, the response values could take negative values, causing the FSI calculation to occasionally yield values outside the theoretical range. Clipping was therefore used to constrain the index within its valid range.

#### Sparseness

The sparseness of neuronal responses to natural image stimuli was quantified using the Hoyer sparseness measure (Hoyer, 2004). For each neuron, responses across repeated presentations of each image were first averaged to obtain a stimulus-response vector:

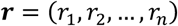

where *n* denotes the number of images, and *r_i_* represents the mean response of the neuron to the *i*-th image across repeated trials. Before calculating sparseness, responses were z-score normalized across stimuli. This normalization was applied to ensure fair comparisons across datasets and data types, particularly because responses in the previously published Utah-array MUA dataset (Papale et al., 2025) were provided in z-scored form. By placing responses from different sources on a common scale, z-score normalization ensured that differences in sparseness primarily reflected the distributional structure of neuronal responses rather than differences in response magnitude or measurement scale.

The response sparseness was then calculated as:

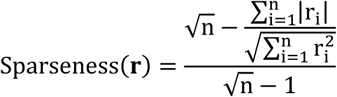

Where 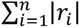 corresponds to the L1 norm of the response vector r, and 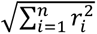 corresponds to its L2 norm. The sparseness value ranges from 0 to 1, with larger values indicating greater stimulus selectivity and responses concentrated on a smaller subset of stimuli.

For the analyses in Fig. 5, we first selected recording sessions containing the shared Random Image Set 1 to ensure consistency of visual stimuli across units. Units were then ranked according to their response sparseness. We selected the 50 units with the highest sparseness values and the 50 units with the lowest sparseness values to form sparse and non-sparse neuronal subsets, respectively.

#### Randomness

To quantify the randomness of neuronal visual responses, we developed a quantitative metric based on spatial autocorrelation analysis. This metric evaluates whether the neural encoding function, conceptualized as a mapping from image feature space to neural response space, preserves local continuity. If a neuron exhibits a continuous encoding function, images that are close in feature space should evoke similar neuronal responses. Conversely, if visually similar stimuli produce dissimilar responses, the neuronal representation is considered more random.

We adopted Moran’s I statistic (Moran, 1950), a classical measure of spatial autocorrelation, to quantify the relationship between stimulus similarity structure and neuronal responses. Moran’s I measures whether observations associated with spatially neighboring locations exhibit correlated values and is defined as:

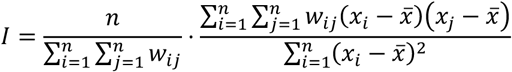

where *n* is the number of spatial units, *x_i_* represents the observation at location *i*, *x̅* denotes the mean of all observations, and *w_ij_* represents the spatial weight between locations *i* and *j*. Higher Moran’s I values indicate stronger spatial clustering, where neighboring locations tend to have more similar values.

To define the image feature space, we extracted high-dimensional image representations using AlexNet (Krizhevsky et al., 2017), a convolutional neural network pretrained on an image classification task. Each stimulus image was passed through AlexNet, and activation maps were extracted from intermediate layers (layer1–layer5, fc6–fc8). These activation maps were flattened into one-dimensional feature vectors, defining the corresponding feature spaces of different network layers.

Based on Moran’s I, we defined a neuronal response randomness metric, where neuronal responses to individual images served as observations and the AlexNet feature space provided the underlying spatial structure. The spatial weight matrix was constructed based on cosine distances between image feature vectors. Because Moran’s I quantifies response continuity rather than randomness, the metric was inverted such that larger values correspond to more random neuronal representations:

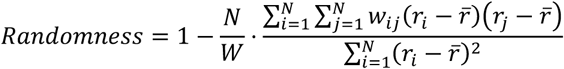

where

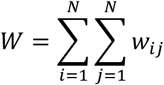

and

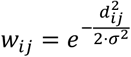

Here, *N* represents the total number of image stimuli, *r_i_* represents the response of a neuron to image *i*, and *d_ij_* denotes the cosine distance between the feature vectors of images *i* and *j* in a given AlexNet layer. The scale parameter *σ* was set to the median pairwise feature distance.

For subsequent statistical analyses, we further transformed the metric as:

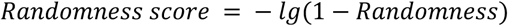

#### Half-peak latency

For each unit, we averaged responses across all natural image stimuli to obtain the stimulus-averaged PSTH. The peak response was identified from this PSTH. The half-peak latency was defined as the earliest time point after stimulus onset at which the response exceeded the average of the baseline firing rate and the peak firing rate.

### Population and dimensionality analyses

For the population analyses presented in Fig. 3, we included only units from the 24 recording sessions that shared the same random image set (Random Image Set 1) to ensure consistency of visual stimuli across sessions.

#### Neural response space

The responses of a neuronal population to a large set of visual stimuli can be represented in a neural response space, where each point corresponds to one sample of the population state evoked by a single image stimulus. The coordinates of each point along individual axes represent the responses of the corresponding neurons, providing a representation of the population activity pattern associated with each stimulus.

To characterize the low-dimensional neural manifold underlying population responses, we applied principal component analysis (PCA) to the visual responses of all n recorded neurons. In an n-dimensional response space, PCA extracts n principal components, each of which represents a linear combination of neuronal firing rates. These components are ordered sequentially to maximize the explained covariance of the population response. The contribution of each principal component was quantified by its explained variance relative to the total variance of the original response matrix.

Fig. 3H shows the cumulative explained variance as a function of the number of retained principal components for three neural populations: SUA from Neuropixels 2.0 (SUA NP2), MUA from Neuropixels 2.0 (MUA NP2), and MUA from Utah arrays (MUA Utah).

#### Effective dimensionality

To estimate the intrinsic dimensionality of neural representations, we used effective dimensionality (ED), a simple linear dimensionality measure (Del Giudice, 2021; Pirkl et al., 2012):

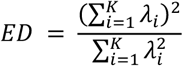

where *λ_i_* represents the *i*-th eigenvalue of the covariance matrix of the population response, and *K* denotes the total number of eigenvalues.

To calculate ED, we constructed a population response matrix consisting of neuronal responses to all visual stimuli, computed the covariance matrix and its eigenvalues, and then estimated ED according to the above definition. To enable fair comparisons across different neural populations, we randomly subsampled an equal number of units from each population before calculating ED.

For Fig. 3I, 100 units were randomly sampled from each neural population (SUA NP2, MUA NP2, and MUA Utah) to estimate ED. For Fig. 5A, 40 units were randomly sampled from sparse and non-sparse neuronal populations to compare their representational dimensionality. This subsampling procedure was repeated multiple times, and the mean ED was used for statistical comparisons.

#### Blockiness

To quantify the modular structure of neuronal response correlation matrices, units were grouped into k functional clusters using hierarchical clustering. The optimal number of clusters was determined by maximizing the silhouette score.

Blockiness was defined as the difference between the mean pairwise correlation coefficient of neuronal pairs within the same cluster and that of neuronal pairs across different clusters:

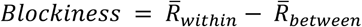

where *R̅_within_* represents the average response correlation between neuron pairs within the same functional cluster, and *R̅_between_* represents the average response correlation between neuron pairs belonging to different clusters.

For Fig. 3G, blockiness was calculated by randomly sampling 100 units from each neural population (SUA NP2, MUA NP2, and MUA Utah).

### Encoding analysis

#### Axis model

We extracted the 4,096-dimensional activations of the FC6 layer of AlexNet for all stimulus images and performed principal component analysis (PCA) to reduce the dimensionality to 50 dimensions, thereby constructing a low-dimensional object space. To compute the preferred axis, defined as the axis capturing the largest variance in image-evoked responses, we used linear regression to estimate the coefficients in the equation *R* = *c* · *F* + *c*_0_, where *R* denotes the response vector of a neuron across the image set, *F* denotes the 50-dimensional object-feature matrix of the image set, and *c*_0_ denotes a constant offset.

For each cell, we estimated the preferred axis using 90% of the images and then projected the remaining 10% of the images onto the resulting axis. This procedure was repeated ten times, with a different subset of images held out as the test set in each iteration. We calculated the correlation between the neural responses and the projection values across all images in the test set. This correlation was normalized by the noise ceiling to obtain the explainable variance, which served as the performance metric.

The mean of the ten estimated preference axes was taken as the overall preference axis of each cell. We then projected all images onto this overall preference axis and divided the resulting projection values into bins. For each bin, we calculated the mean neural response across all images within that bin. These binned responses were used to visualize the explanatory power of the axis encoding.

#### Deep neural network feature transfer model for predicting neuronal responses

Stimulus images were preprocessed before being used as model inputs. The original images had a resolution of 330×330 pixels. Before being fed into the model, images were center-cropped to retain the middle 256×256 pixels and subsequently resized to 200×200 pixels using bicubic interpolation while preserving the aspect ratio.

The feature transfer model consisted of a feature extractor followed by a two-layer perceptron. Image features were extracted from ResNet-50 (He et al., 2016) using the output of layer 153 (conv5_block1_add_bn). For each stimulus image, this layer generated a spatial feature map with dimensions 7×7×2048. Higher layers of DNN have been shown to better predict responses in the inferotemporal (IT) cortex (Schrimpf et al., 2020; Yamins et al., 2014). We evaluated multiple ResNet-50 layers and selected layer 153 because it yielded the highest prediction accuracy for neuronal responses (Fig. S5).

We used the Keras implementation of ResNet-50 with weights pretrained on the ImageNet classification task. The extracted image features were then mapped onto neuronal responses using a two-layer perceptron. The hidden layer consists of 400 units, which was empirically determined to provide improved prediction performance. An exponential linear unit (ELU) nonlinearity was applied to the hidden layer to enhance the representational capacity of the model.

To improve generalization and reduce overfitting, dropout regularization was applied in the model. During forward propagation, a fraction of hidden units was randomly deactivated, encouraging the model to learn more robust representations. Hyperparameter optimization indicated that a dropout rate of 0.5 provided good performance. In addition, L1 regularization was applied to the connection weights of the perceptron. L1 regularization promotes sparse weight distributions, encouraging the selection of more relevant and informative features, thereby reducing the risk of overfitting and improving generalization performance.

Before DNN training, neuronal firing rates were scaled by a factor of 0.1 to align the numerical range of neural responses with that of the model outputs. All models were optimized using stochastic gradient descent with the Adam optimizer. The batch size was set to 20. Stimulus images were randomly divided into training, validation, and test sets with an 8:1:1 ratio.

During model fitting, network parameters were optimized to minimize the mean squared error (MSE) between predicted and measured neuronal responses on the training set. To prevent overfitting and ensure optimal generalization, early stopping was implemented based on the MSE between predicted and measured responses on the validation set. Training was terminated if the validation MSE did not improve for 50 consecutive epochs, or after a maximum of 80 epochs. The model achieving the best validation performance during training was saved as the final model. This procedure allowed us to capture optimal model performance while avoiding unnecessary training iterations.

Model performance was evaluated by calculating the Pearson correlation between predicted and measured neuronal responses to images in the test set. Because trial-to-trial noise and intrinsic response variability limit the maximum correlation that a noise-free model can achieve with the measured responses, we estimated the noise ceiling using split-half reliability measured on the test set (see Methods: Reliability). The split-half reliability was Spearman–Brown corrected to estimate the reliability of the full dataset. Because split-half reliability represents the proportion of explainable response variance rather than the maximum achievable correlation, the corresponding noise ceiling for a correlation-based performance metric was defined as the square root of the reliability. The prediction correlation of each model was therefore divided by the square root of the reliability to obtain the noise-normalized performance, which was used as the final performance metric.

### Decoding analysis

For the decoding analysis, sparse/non-sparse and random/non-random neuronal subsets in recording sessions containing the shared Random Image Set 1 were used.

We evaluated population coding performance in two visual tasks: categorization and identification. In the categorization task, stimulus labels indicated whether each image belonged to the face or non-face category, with 50 face images and 50 non-face images included. In the identification task, labels represented individual image identities. The stimulus set for this task was generated by collecting each unit’s preference images and removing duplicate images, resulting in a total of 79 unique images for sparse/non-sparse and 86 for random/non-random.

Both categorization and identification tasks were decoded using linear classifiers. For the categorization task, the classifier discriminated between face and non-face images. For the identification task, the classifier predicted the corresponding image identity represented as a one-hot encoded output vector.

The dataset was divided into training and testing sets. The training set consisted of five repetitions of each image, whereas the testing set consisted of the remaining single repetition. During classifier training, different levels of L1 regularization were applied to progressively enforce sparsity in decoder weights. As the strength of L1 regularization increased, the number of non-zero input weights contributing to each output node gradually decreased. When the number of non-zero input weights for each output node was reduced to k or fewer, the corresponding decoding accuracy was recorded. By progressively increasing L1 regularization strength, k was varied from 16 to 1, allowing us to characterize the relationship between decoding performance and the number of units contributing to the decoder.

### Comparative Analysis of Artificial and Biological Neural Representations

To characterize the response properties of artificial neurons and compare them with those of biological neurons, we treated individual units in ResNet-50 as artificial neurons and measured their responses to the same natural images presented to the monkeys. Images were resized to 224 × 224 pixels and processed using the standard Keras preprocessing pipeline before being input to an ImageNet-pretrained ResNet-50. We extracted activations from the conv5_block1_out layer (layer 154), which produced a 7 × 7 × 2,048 activation map for each image, treating each spatial position within each channel as a candidate artificial neuron. We randomly sampled 10,000 candidate units and retained those whose maximum activation across the stimulus set exceeded 10, resulting in approximately 5,000 artificial neurons. For each artificial neuron, the activation elicited by each image was taken as its visual response.

We quantified two response properties using the same measures applied to the biological neurons: feature randomness and lifetime sparseness. Feature randomness was quantified using Moran’s I based on the feature-space relationships between stimulus images, with image features defined by layer 5 of AlexNet. Lifetime sparseness was quantified using the Hoyer sparseness index based on the distribution of each artificial neuron’s responses across images. These measures were used to compare the distributions of feature randomness and lifetime sparseness between artificial and biological neurons.

For the decoding analysis, we followed the same procedure described above for biological neurons, with the following modifications. Artificial neurons were divided into sparse/non-sparse and random/non-random populations by randomly selecting units from the top and bottom 5% of the corresponding lifetime sparseness and feature randomness distributions, respectively. To simulate trial-to-trial variability, independent Gaussian noise (*σ* = 1.0) was added to the mean response of each artificial neuron for each image, generating six simulated trials per image. The same categorization and identification tasks were then evaluated using linear decoders. For categorization, the stimulus set comprised 50 face and 50 non-face images. For identification, preference images from the selected artificial neurons were pooled, duplicate images were removed, and 100 images were randomly selected. All other aspects of the decoding procedure, including the training/testing scheme, L1 regularization, and evaluation of decoding accuracy as a function of the number of effective connections, were identical to those described above.

To examine how artificial units with different levels of feature randomness and response sparseness contributed to the network’s own classification performance, we evaluated ResNet-50 units on the Random Image Set I using a binary face/non-face classification task. For each image, the activation of each unit in the final feature-extraction layer of ResNet-50 was taken as the unit’s response. Feature randomness and lifetime sparseness were quantified for each unit using the same measures described above.

To assess the contribution of feature randomness, units were ranked according to their feature-randomness values and divided into five equal-sized quintiles. For each quintile, we retained only the units within that quintile and evaluated the network’s face/non-face classification accuracy using the corresponding network outputs. The same procedure was applied to lifetime sparseness, with units ranked according to their sparseness values and classification accuracy evaluated separately for each of the five quintiles. Thus, each analysis assessed the classification performance supported by a distinct 20% subset of units spanning the corresponding range of feature randomness or response sparseness.

### Comparison of Neural Coding Schemes

To compare different coding schemes, we contrasted the measured SUA responses with responses reconstructed using the axis model. The axis model constrained each neuron’s response to the component that could be explained by the 50-dimensional feature axes described above, thereby removing response components not captured by the axis-based representation. For each neuron and image, the axis-fitted response was obtained from the model-predicted response based on its projection onto the 50 feature axes. Thus, the measured and axis-fitted populations had identical dimensions, while differing only in the extent to which their responses were constrained by the low-dimensional axis representation.

We evaluated the two response populations using an identical image-identification decoding procedure. For each neuron, responses were z-scored across images. The analysis was repeated 10 times independently. In each repetition, 40 images were randomly selected, and independent Gaussian noise (*σ* = 1.0) was added to the mean population response of each image to generate five training and five testing simulated trials. The measured and axis-fitted populations used the same images and were applied the same noise, ensuring that differences in decoding performance reflected differences in the underlying population response geometry rather than differences in stimulus or noise sampling.

Image identity was decoded using a one-versus-rest linear SVM with L1 regularization. We systematically varied the regularization parameter *C* and evaluated decoding performance on the independent simulated test trials using 40-way classification (chance level = 2.5%). Connection density was defined as the proportion of non-zero weights among all potential connections. For each value of *C*, test accuracy and the resulting connection density were calculated separately and then averaged across the 10 repetitions, with the standard error reported across repetitions.

Since the same value of *C* could produce different numbers of nonzero connections in the measured and axis-fitted populations, decoding performance was compared as a function of the actual connection density rather than the regularization parameter. This analysis quantified the complexity of sparse linear readout required by each population representation to achieve a given level of image-identification performance, under matched synthetic trial variability and an identical linear decoding framework.

## Data and code availability

The dataset will be available upon publication. Image stimuli were derived from the ImageNet (Deng et al., 2009) database (https://image-net.org/downloadimages.php). Custom code for data preprocessing, model training and related analysis will be available upon publication.

## Acknowledgements

We thank the Peking University Laboratory Animal Center for excellent animal care. This work was supported by Brain Science and Brain-like Intelligence Technology --National Science and Technology Major Project, grant no.2022ZD0204600 (to S.T.) and funds from the Peking-Tsinghua Center for Life Sciences (to S.T.).

## Supplementary Figures

**Fig. S1.**
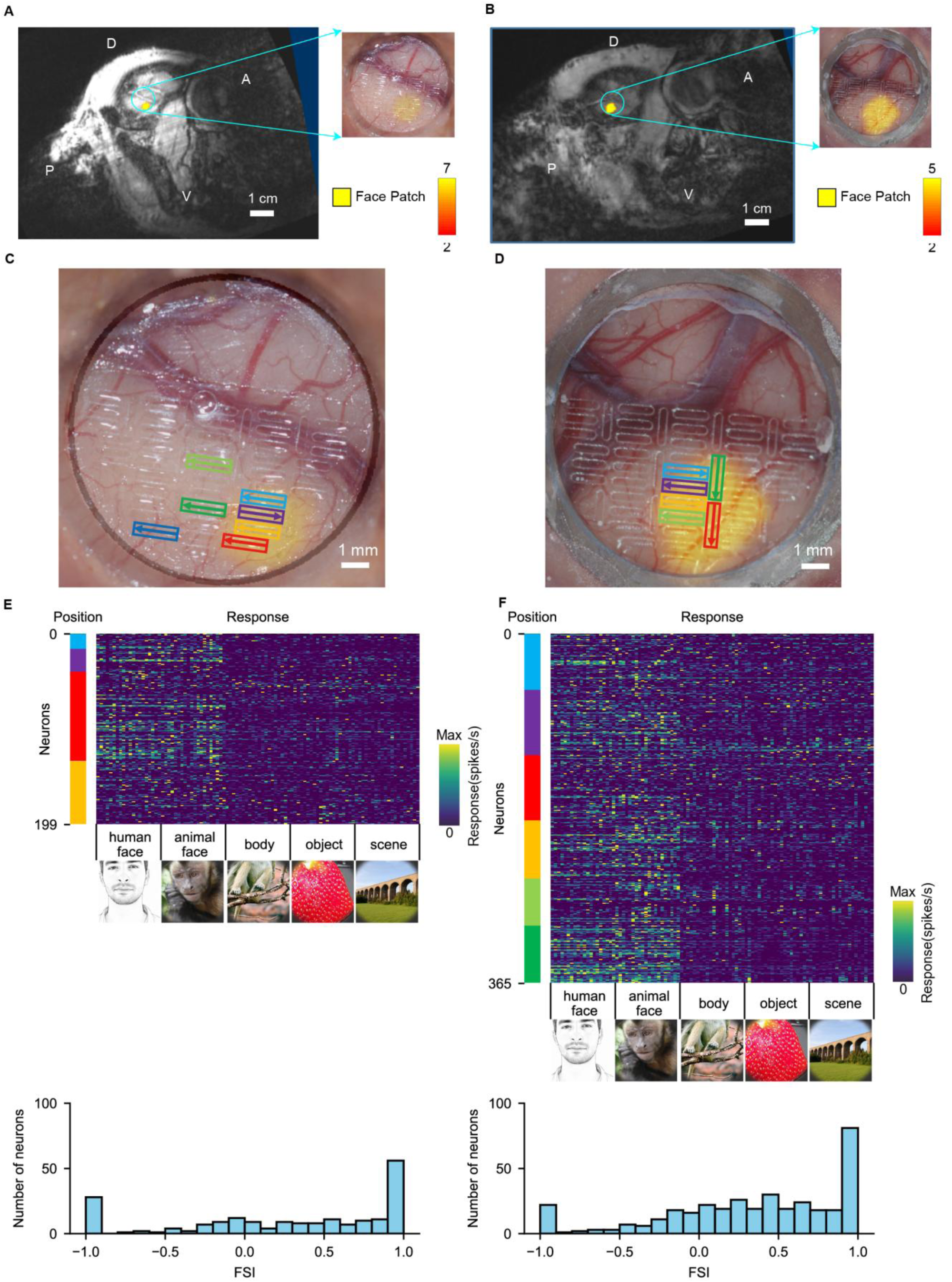
Face-patch localization, recording sites and neuronal selectivity. **A, B,** Localization of the face patch in two macaques and projection of its position onto the chamber window. The fMRI-defined face patch (yellow) was aligned to the chamber view using landmark surface blood vessels visible in both the MR images and the chamber photographs. D, dorsal; V, ventral; A, anterior; P, posterior. **C, D,** Neuropixels recording locations in the two macaques. Yellow shading indicates the fMRI-defined face patch. Each outlined rectangle denotes one recording site and probe orientation, with the arrow pointing from shank 0 toward shank 3. Outline colors correspond to the recording sessions listed in Table 1. **E, F,** Localizer response profiles and face-selectivity-index (FSI) distributions for the recorded neurons in the two macaques (*n* = 199 and 365 neurons, respectively). In each response matrix, rows represent neurons and columns represent localizer images, grouped into human-face, animal-face, body, object and scene categories. Human face image example has been replaced with cartoon illustrations. The colored strip beside each matrix indicates the recording location using the same color scheme as in **C** and **D**. Histograms below show the corresponding distributions of FSI values.

**Table 1.** Experimental recording sessions. **Session id:** recording session number; **Monkey:** subject identifier; **Position:** electrode implantation location, with colors corresponding to those in **Fig. S1**; **In face patch**: indicates whether the recording was within the face patch; **imec id**: electrode identifier; **Random imgset choice**: the image set used (two sets available); **Date**: recording date.

| Session id | Monkey | Position | In face patch | imec id | Random imgset choice | Number of images | Date |
| --- | --- | --- | --- | --- | --- | --- | --- |
| 1 | A | 1 | Yes | 0 | 1 | 3000 | 20250324 |
| 2 | A | 2 | Yes | 1 | 1 | 3000 | 20250324 |
| 3 | A | 3 | Yes | 0 | 1 | 3000 | 20250326 |
| 4 | A | 3 | Yes | 0 | 2 | 3000 | 20250327 |
| 5 | A | 4 | Yes | 0 | 1 | 3000 | 20250407 |
| 6 | A | 4 | Yes | 0 | 1 | 3000 | 20250408 |
| 7 | A | 3 | Yes | 0 | 1 | 3000 | 20250411 |
| 8 | A | 3 | Yes | 0 | 1 | 3000 | 20250412 |
| 9 | A | 5 | No | 0 | 1 | 3000 | 20250414 |
| 10 | A | 5 | No | 0 | 2 | 3000 | 20250415 |
| 11 | A | 6 | No | 0 | 1 | 3000 | 20250417 |
| 12 | A | 7 | No | 0 | 1 | 3000 | 20250418 |
| 13 | B | 1 | Yes | 0 | 1 | 3000 | 20251008 |
| 14 | B | 2 | Yes | 1 | 1 | 3000 | 20251008 |
| 15 | B | 1 | Yes | 0 | 1 | 2400 | 20251009 |
| 16 | B | 2 | Yes | 1 | 1 | 2400 | 20251009 |
| 17 | B | 3 | Yes | 0 | 1 | 2400 | 20251011 |
| 18 | B | 3 | Yes | 0 | 1 | 3000 | 20251012 |
| 19 | B | 4 | Yes | 0 | 1 | 3000 | 20251014 |
| 20 | B | 5 | Yes | 1 | 1 | 3000 | 20251014 |
| 21 | B | 4 | Yes | 0 | 1 | 3000 | 20251015 |
| 22 | B | 5 | Yes | 1 | 1 | 3000 | 20251015 |
| 23 | B | 4 | Yes | 0 | 2 | 2400 | 20251016 |
| 24 | B | 5 | Yes | 1 | 2 | 2400 | 20251016 |
| 25 | B | 4 | Yes | 0 | 1 | 2400 | 20251017 |
| 26 | B | 5 | Yes | 1 | 1 | 2400 | 20251017 |
| 27 | B | 6 | Yes | 0 | 1 | 2400 | 20251019 |
| 28 | B | 6 | Yes | 0 | 1 | 3000 | 20251020 |

**Fig. S2.**
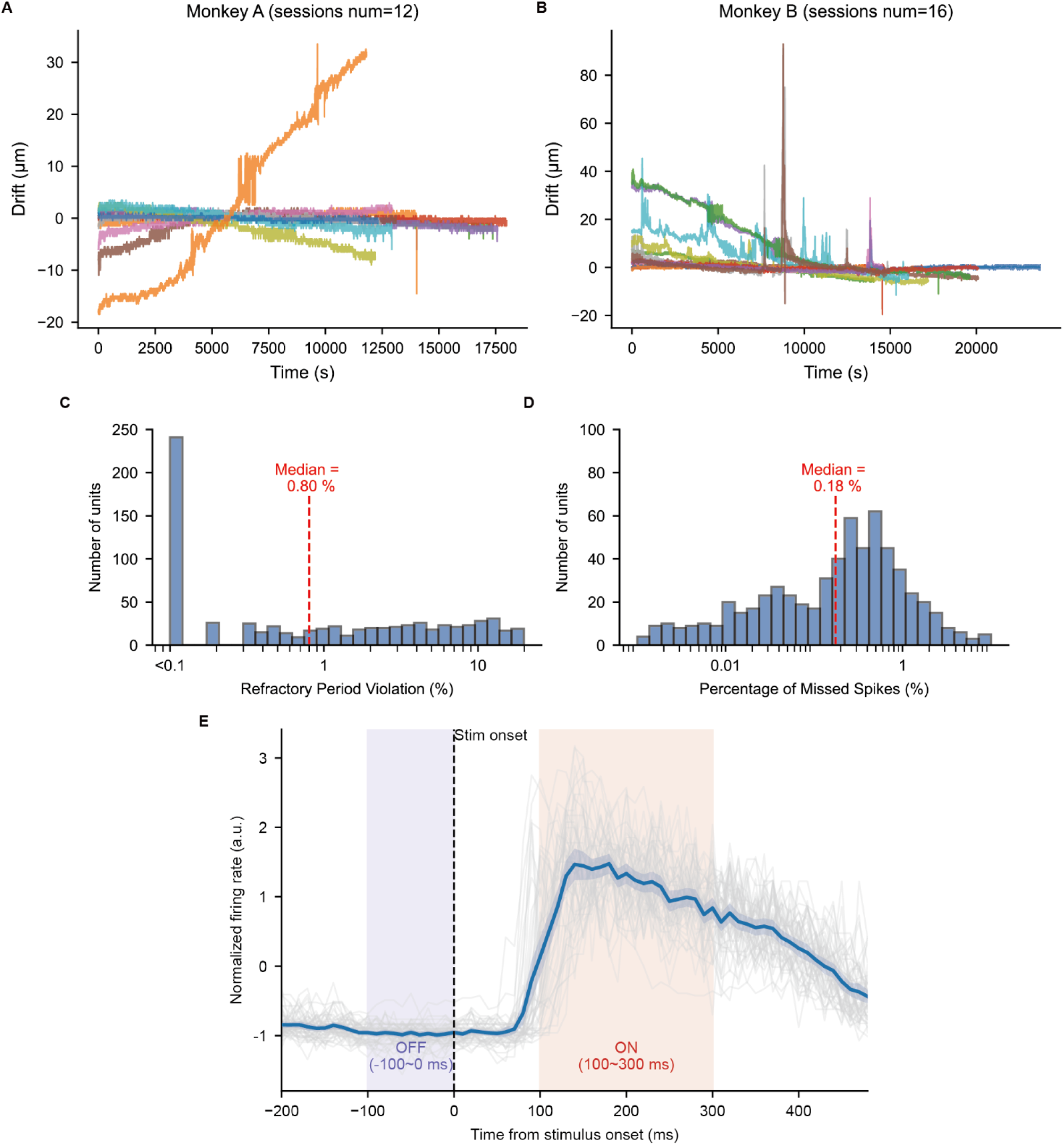
Recording stability, spike-sorting quality, and stimulus-evoked response dynamics. **A, B,** Estimated probe–tissue drift over recording time for Monkey A (**A**; 12 sessions) and Monkey B (**B**; 16 sessions). Drift was estimated by batch correlation using 2-s windows, and each colored trace represents one recording session. **C,** Distribution of refractory-period violation rates across retained single units. **D,** Distribution of the estimated percentage of missed spikes across retained single units. In **C** and **D**, red dashed lines indicate the medians. **E,** Stimulus-aligned response time courses of 50 randomly sampled single units. Responses were z-scored separately for each unit. Gray curves show individual units, the blue line shows their mean response, and the blue shaded region indicates the 95% confidence interval across units. The vertical dashed line marks stimulus onset. Purple and red shaded regions indicate the baseline window (−100–0 ms) and response window (100–300 ms), respectively, used to quantify visually evoked responses.

**Fig. S3.**
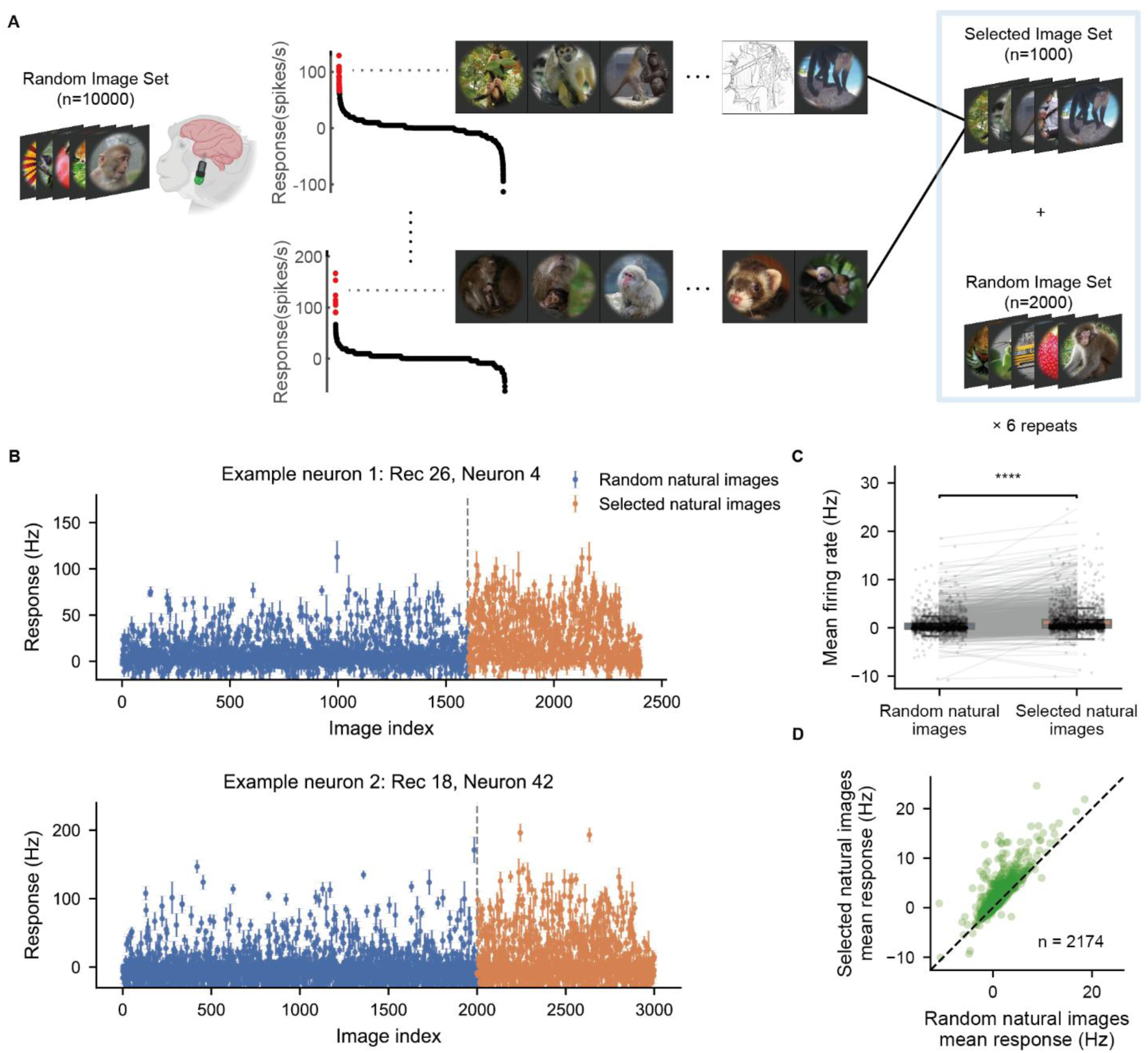
Stimulus-selection procedure and response enrichment in the selected image set. **A,** Schematic of the stimulus-selection procedure. Monkeys first viewed a screening set of 10,000 randomly sampled natural images, each presented once. After offline spike sorting, we selected 1,000 images that elicited strong responses across the identified candidate units and combined them with 2,000 randomly sampled images to form a final set of 3,000 images. We presented each image in the final set six times. Human face image has been replaced with cartoon illustrations. **B,** Responses of two example single units to images in the random and selected sets. Points and error bars indicate the mean ± s.e.m. response across repeated presentations of each image. The vertical dashed line separates images from the two sets. **C, D,** Comparison of each unit’s mean response to images in the random and selected sets. In **C**, points represent individual units, gray lines connect paired measurements from the same unit, and box plots indicate the median and interquartile range. **D** shows the same data as a scatter plot, with each point representing one unit and the dashed line indicating equality between the two conditions. Responses to selected images were significantly higher than responses to random images (two-sided Wilcoxon signed-rank test, *P* < 0.0001, *n* = 2,174).

**Fig. S4.**
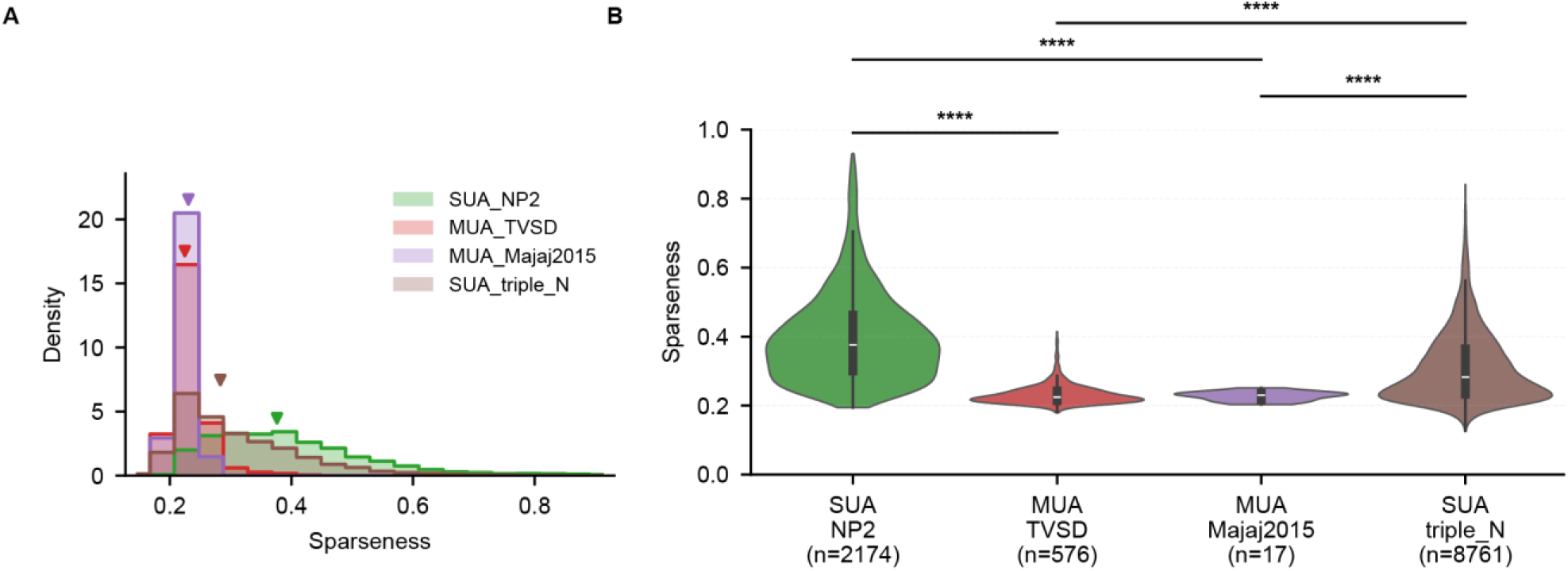
Comparison of response sparseness across Neuropixels SUA and Utah-array MUA datasets. **A,** Density-normalized distributions of response sparseness for NP2 single-unit activity (SUA; present study), Triple-N SUA, TVSD multi-unit activity (MUA), and Majaj et al. (2015) MUA. NP2 and Triple-N SUA were recorded with Neuropixels probes, whereas TVSD and Majaj et al. (2015) MUA were recorded with Utah arrays. Sparseness was computed using the same metric for all four datasets. Arrowheads indicate medians. **B,** Violin plots comparing response sparseness across four data sets. Black boxes indicate the median and interquartile range. Comparisons indicated by horizontal bars were performed using two-sided Mann–Whitney *U*-tests. \**P* < 0.05, \*\**P* < 0.01, \*\*\**P* < 0.001, and \*\*\*\**P* < 0.0001.

**Fig. S5.**
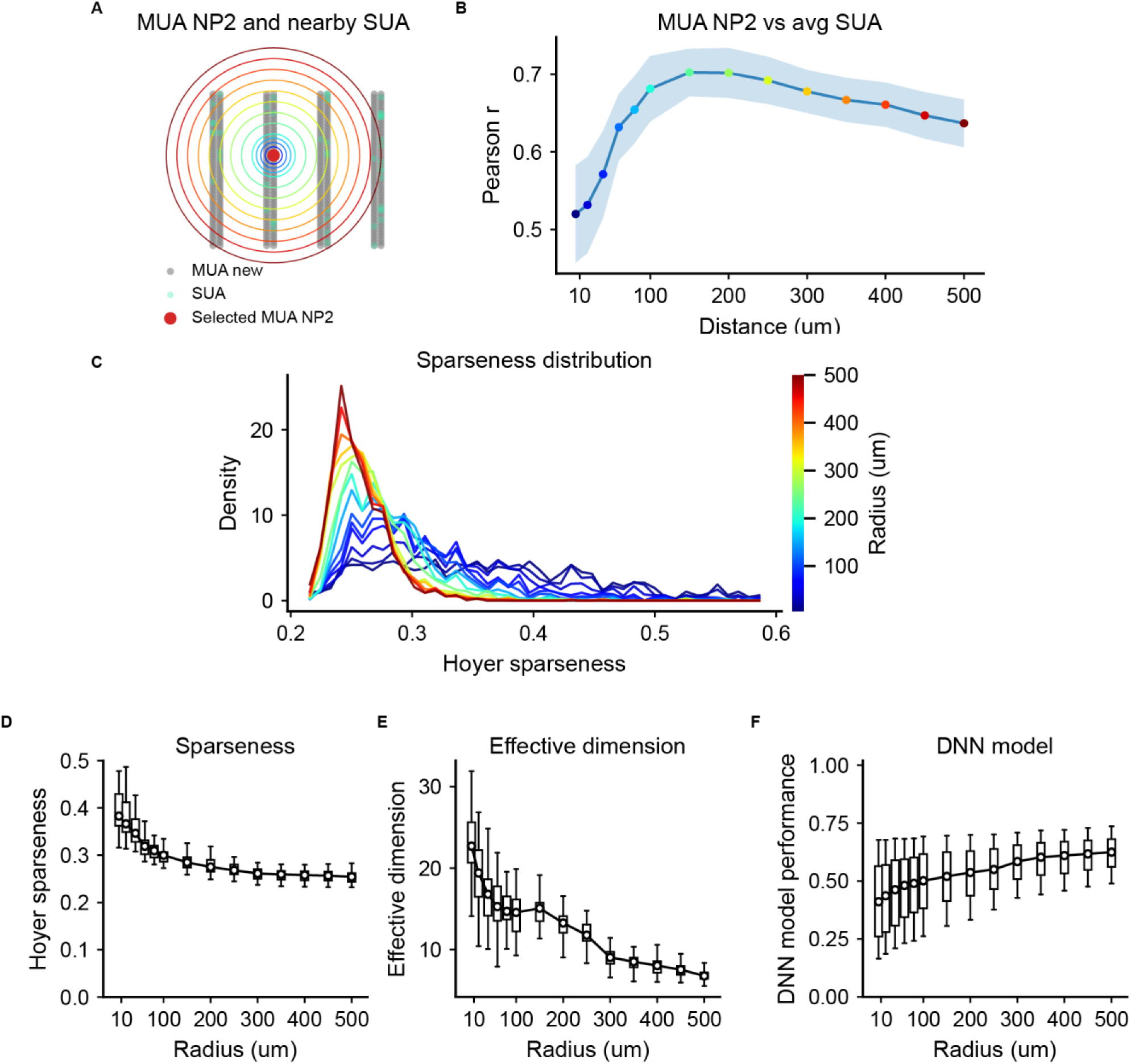
Spatial averaging of SUA reproduces MUA-like response properties. **A,** Schematic of the spatial-averaging analysis. The red point marks a target NP2 MUA channel, green points mark simultaneously recorded SUA, and colored circles indicate pooling radii centered on the target channel. SUA responses within each radius were averaged to generate a simulated MUA response. **B,** Pearson correlation between the measured MUA response profile and the corresponding spatially averaged SUA response profile as a function of pooling radius. The line shows the mean across 30 random samples of MUA sites, and shading indicates the 95% confidence interval. **C,** Density distributions of Hoyer sparseness for spatially averaged SUA responses at each pooling radius; colors correspond to those in **A**. **D-F,** Response sparseness (**D**), effective dimensionality (**E**), and noise-ceiling-normalized DNN prediction performance (**F**) as a function of pooling radius. Box plots summarize the distributions across random samples.

**Fig. S6.**
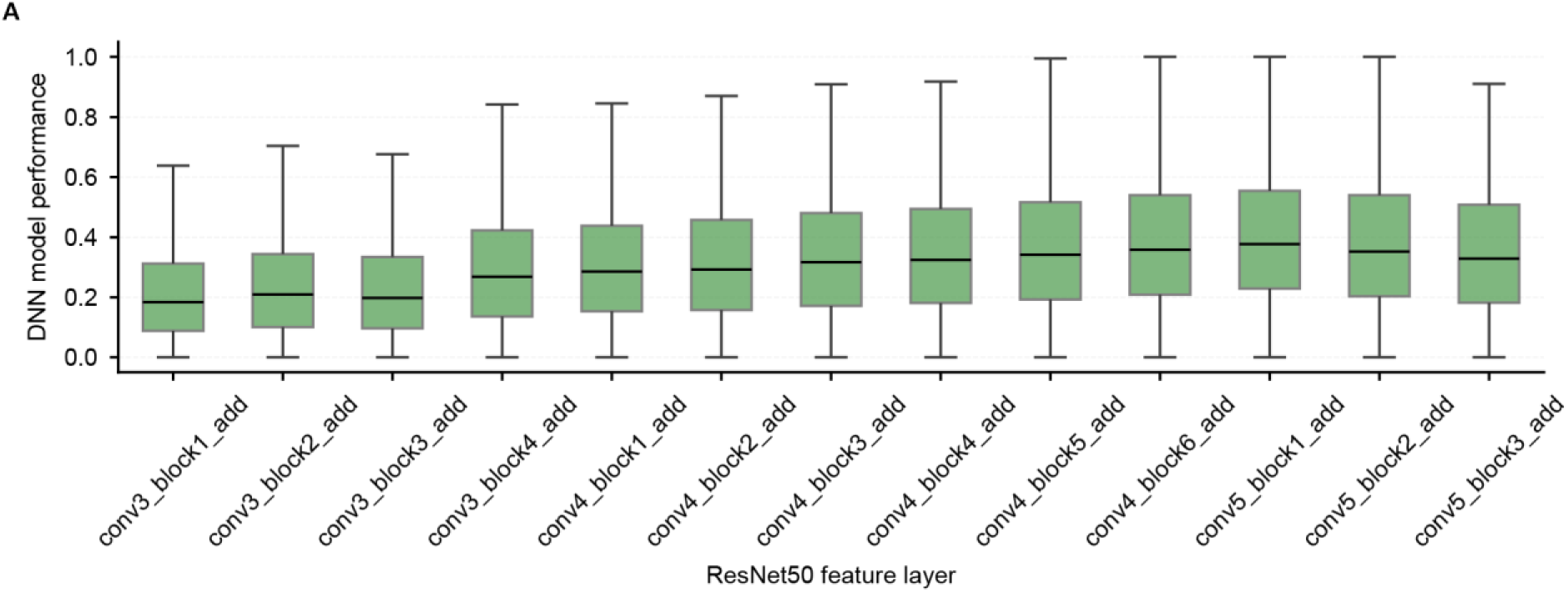
DNN prediction performance across ResNet-50 feature layers. **A,** Noise-ceiling-normalized prediction performance of encoding models constructed from successive ResNet-50 feature layers. Box plots show the distributions across single units; black center lines and boxes indicate the median and interquartile range, respectively. The best-performing layer, conv5_block1_add_bn, was used in subsequent analyses.

**Fig. S7.**
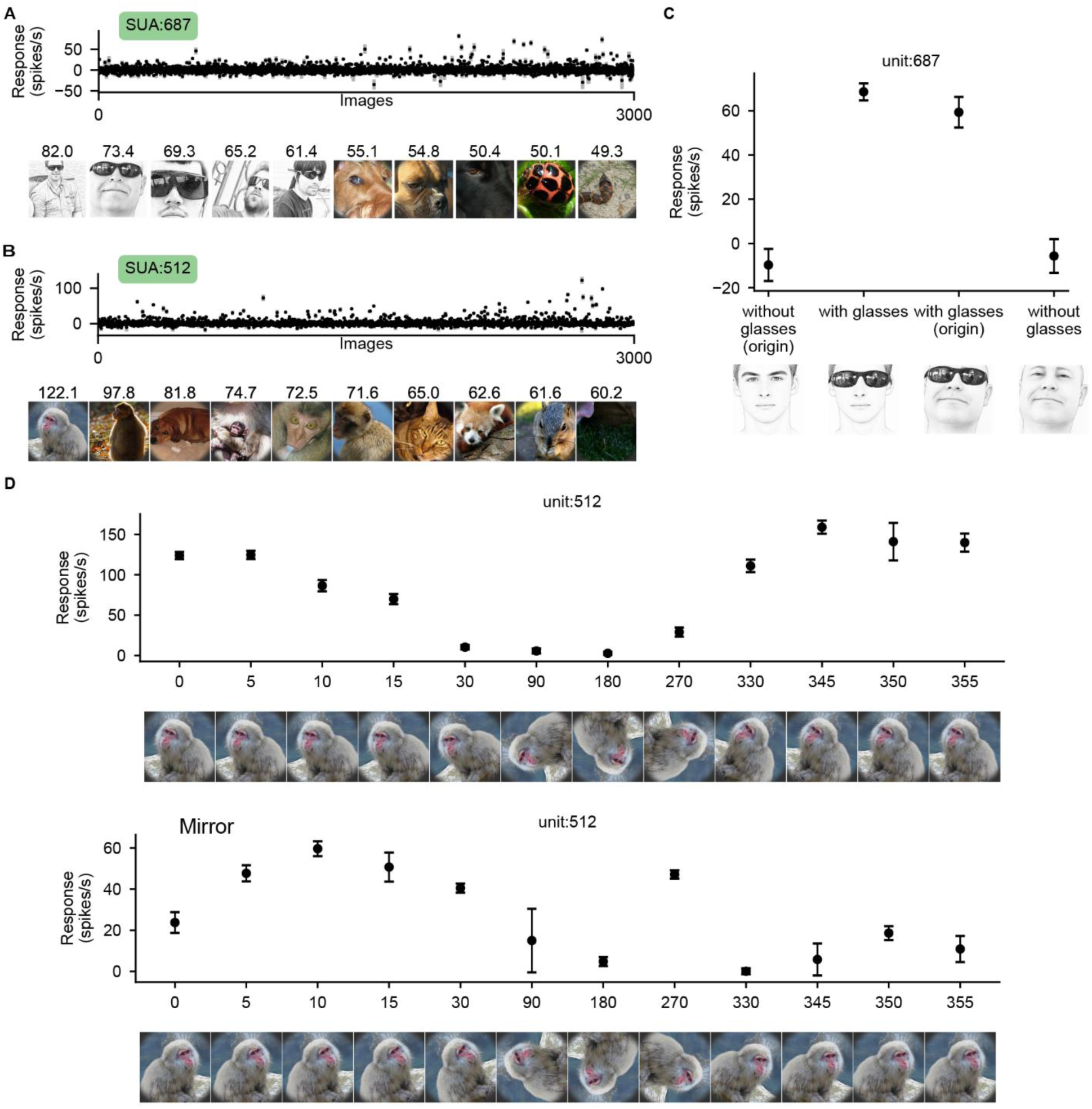
Example single units selective for eyeglasses and face orientation. **A, B,** Response profiles of unit 687 (**A**) and unit 512 (**B**) across 3,000 natural images, together with their ten most effective images. **C,** Responses of unit 687 to original faces with or without eyeglasses and their edited counterparts. Adding eyeglasses to a face without them increased the response, whereas removing eyeglasses from a face that originally contained them reduced the response. **D,** Responses of unit 512 to the original (top) and mirror-flipped (bottom) versions of the same face across in-plane rotation angles. Points and error bars indicate the mean ± s.e.m. across repeated presentations; values above the preferred images in **A** and **B** indicate mean firing rates in spikes/s. For image examples in **A** and **C**, human face images have been replaced with cartoon illustrations.

**Fig. S8.**
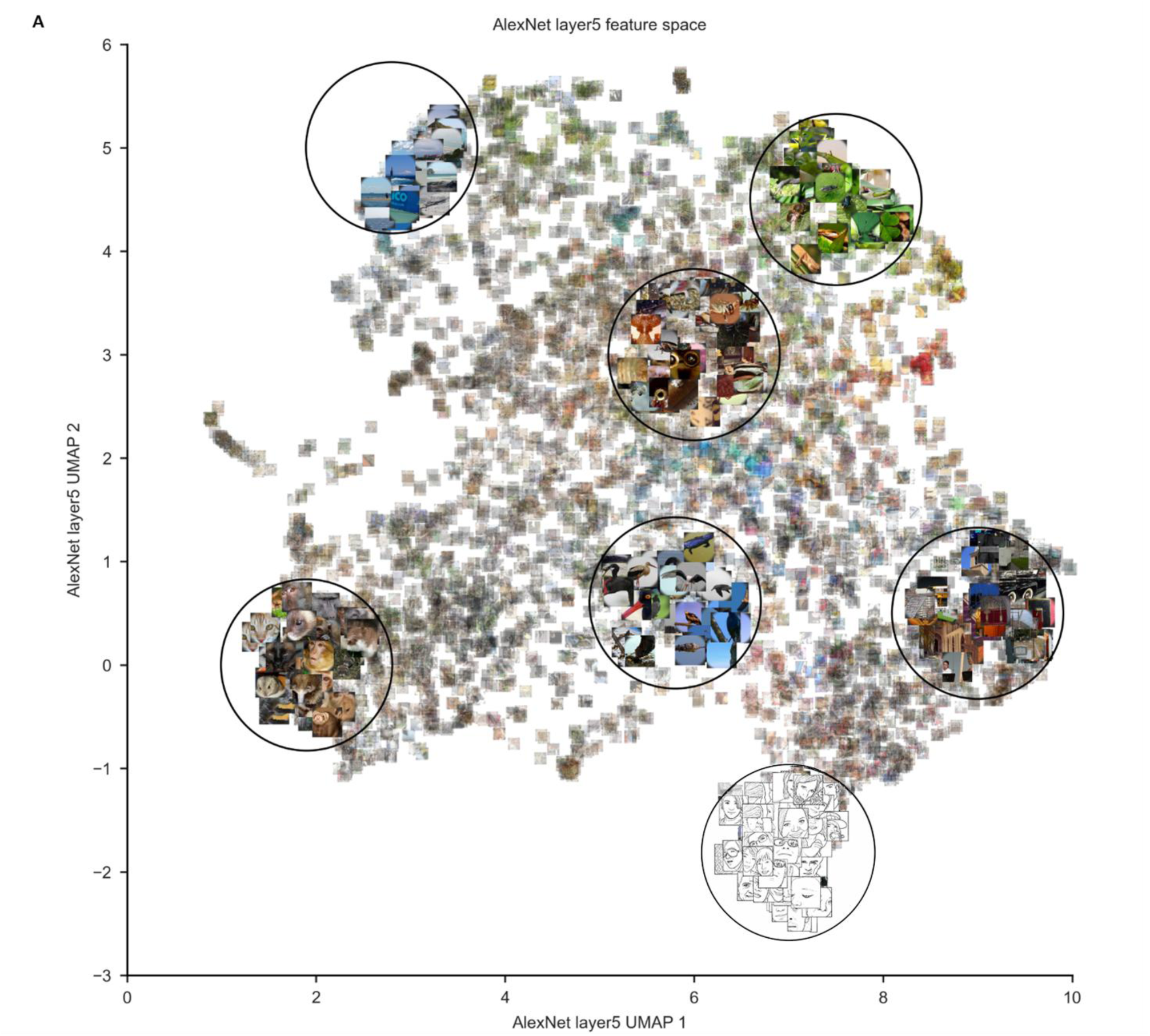
UMAP visualization of the AlexNet layer 5 image-feature space. **A,** Two-dimensional UMAP embedding of the natural images based on features extracted from AlexNet layer 5. Images are displayed at their corresponding UMAP coordinates. Circled regions illustrate local clustering by visual content, including faces, other animals, vegetation, and scenes. Human face images have been replaced with cartoon illustrations.

**Fig. S9A.**
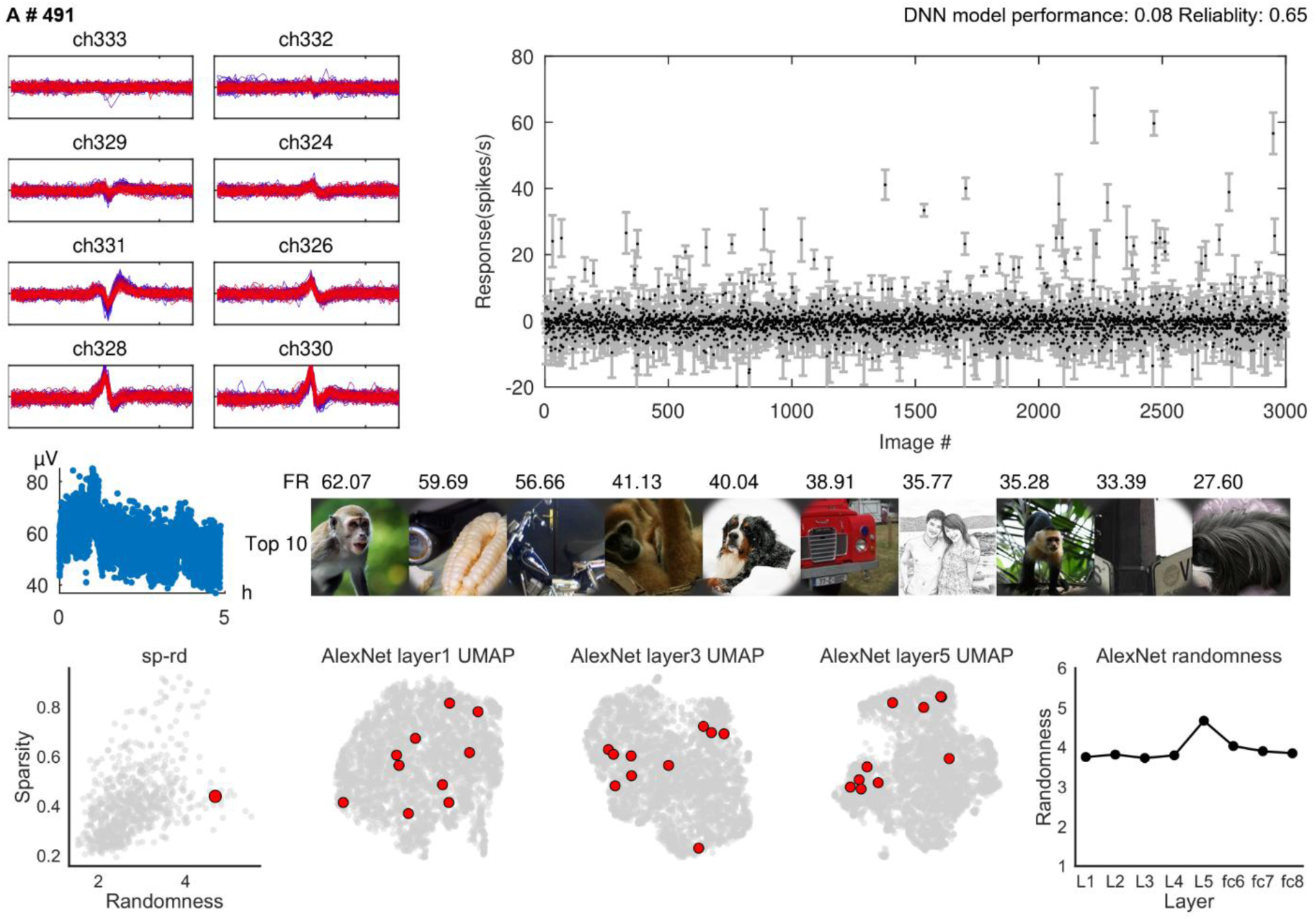
Detailed response properties of an example SUA #491. Multichannel spike waveforms (upper left), mean responses across 3,000 images (upper right), spike amplitude over recording time (middle left), and the ten most effective images with their mean firing rates (middle right). Human face images have been replaced with cartoon illustrations. DNN prediction performance and response reliability are reported above the response profile. The bottom row shows the unit’s position in the joint sparseness-randomness distribution, the locations of its preferred images in UMAP embeddings derived from AlexNet layers 1, 3, and 5, and its feature-randomness values across AlexNet layers. Response points and error bars indicate the mean ± s.e.m. across repeated image presentations. Unit #491 was recorded from Monkey A, session 4.

**Fig. S9B.**
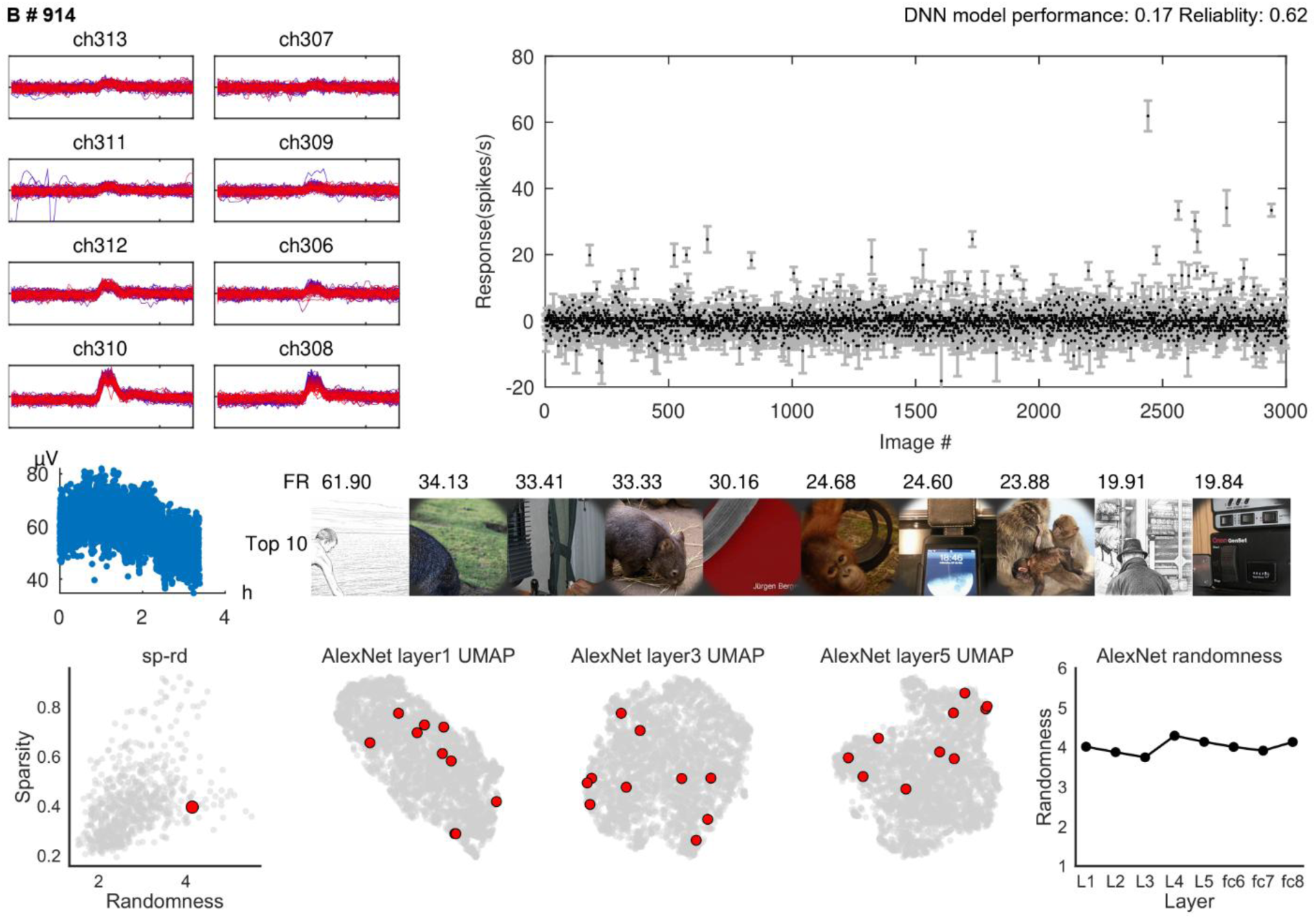
Detailed response properties of example SUA #914. Multichannel spike waveforms (upper left), mean responses across 3,000 images (upper right), spike amplitude over recording time (middle left), and the ten most effective images with their mean firing rates (middle right). Human face images have been replaced with cartoon illustrations. DNN prediction performance and response reliability are reported above the response profile. The bottom row shows the unit’s position in the joint sparseness-randomness distribution, the locations of its preferred images in UMAP embeddings derived from AlexNet layers 1, 3, and 5, and its feature-randomness values across AlexNet layers. Response points and error bars indicate the mean ± s.e.m. across repeated image presentations. Unit #914 was recorded from Monkey A, session 8.

**Fig. S9C.**
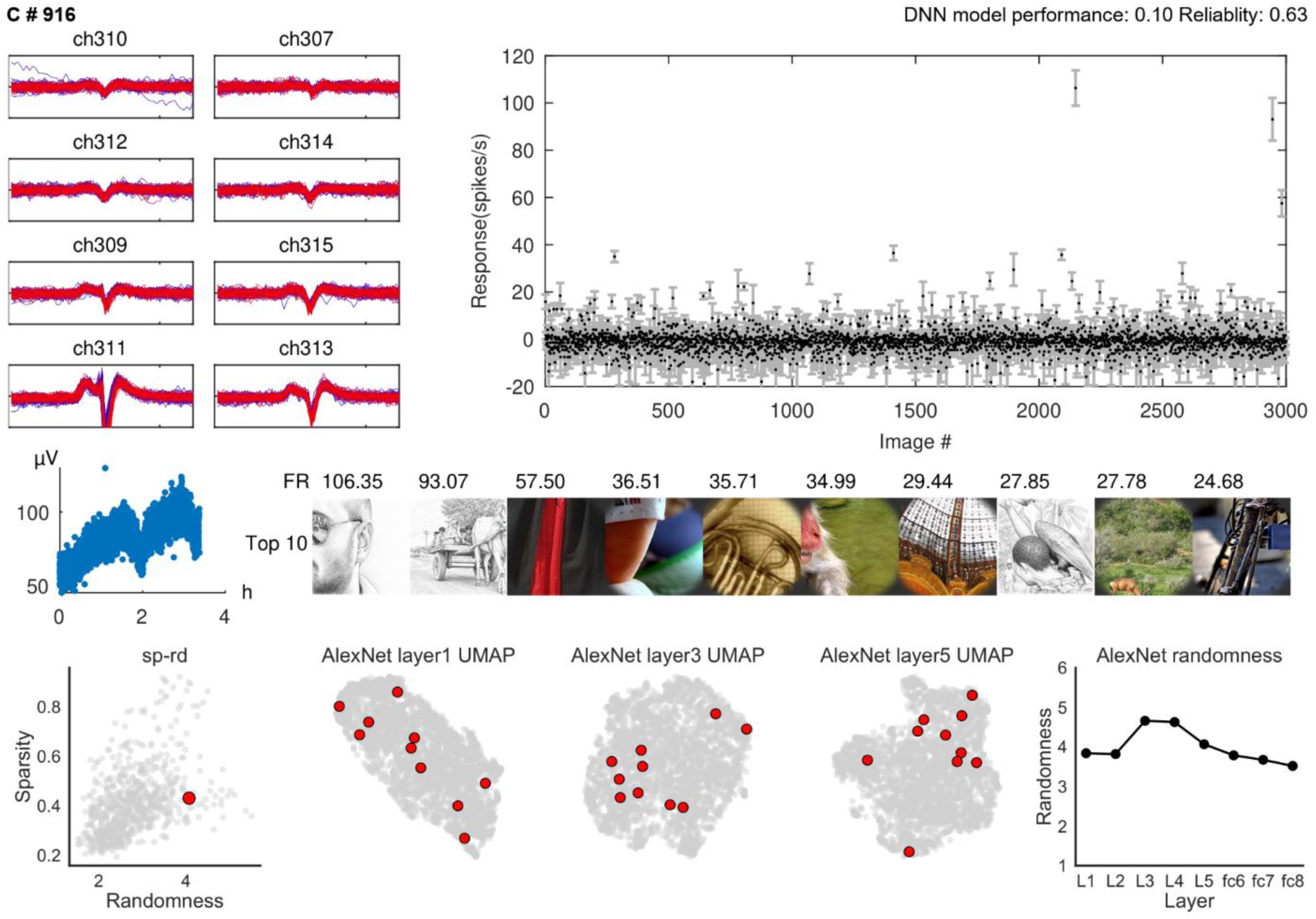
Detailed response properties of example SUA #916. Multichannel spike waveforms (upper left), mean responses across 3,000 images (upper right), spike amplitude over recording time (middle left), and the ten most effective images with their mean firing rates (middle right). Human face images have been replaced with cartoon illustrations. DNN prediction performance and response reliability are reported above the response profile. The bottom row shows the unit’s position in the joint sparseness-randomness distribution, the locations of its preferred images in UMAP embeddings derived from AlexNet layers 1, 3, and 5, and its feature-randomness values across AlexNet layers. Response points and error bars indicate the mean ± s.e.m. across repeated image presentations. Unit #916 was recorded from Monkey A, session 8.

**Fig. S9D.**
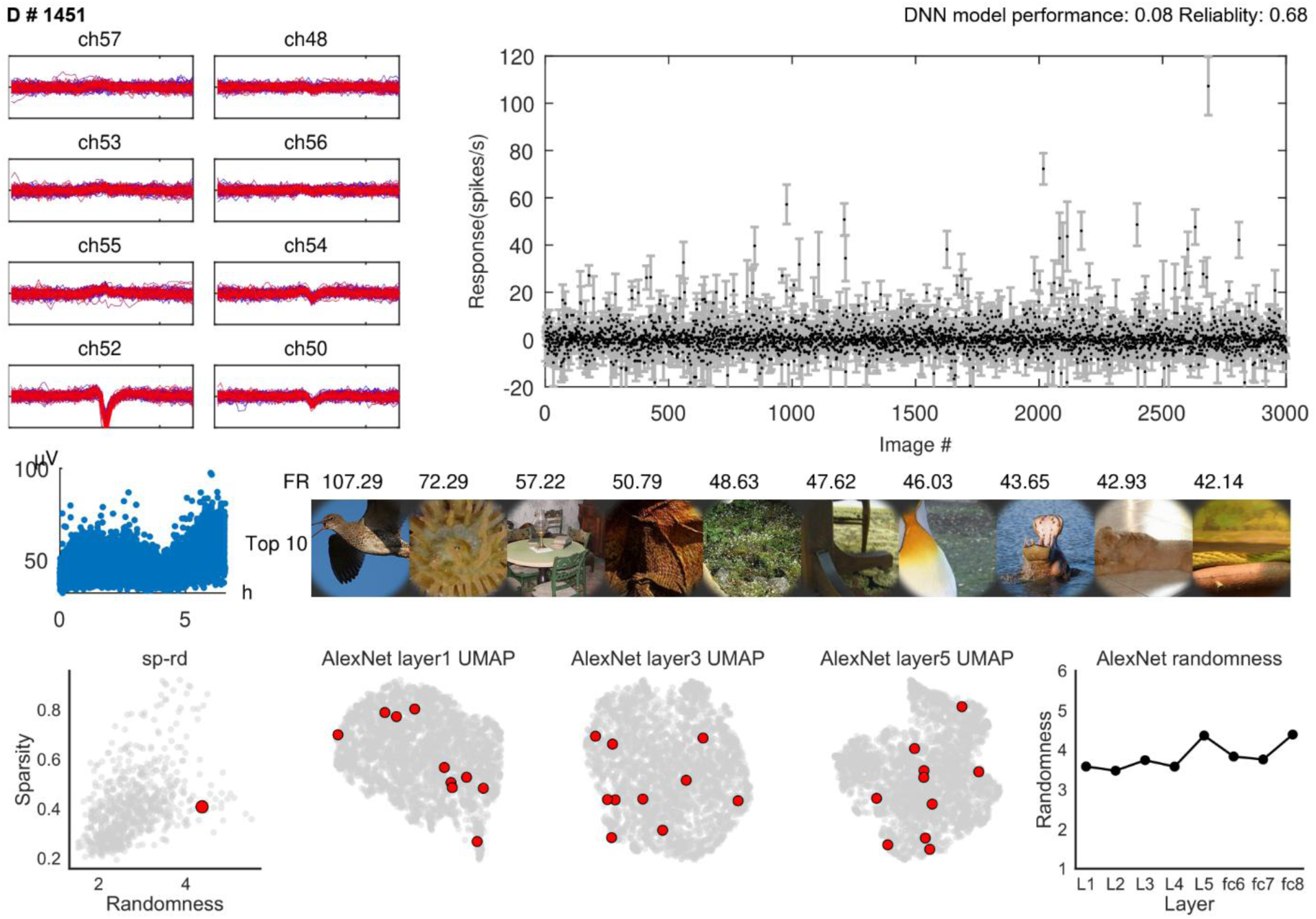
Detailed response properties of example SUA #1451. Multichannel spike waveforms (upper left), mean responses across 3,000 images (upper right), spike amplitude over recording time (middle left), and the ten most effective images with their mean firing rates (middle right). Human face images have been replaced with cartoon illustrations. DNN prediction performance and response reliability are reported above the response profile. The bottom row shows the unit’s position in the joint sparseness-randomness distribution, the locations of its preferred images in UMAP embeddings derived from AlexNet layers 1, 3, and 5, and its feature-randomness values across AlexNet layers. Response points and error bars indicate the mean ± s.e.m. across repeated image presentations. Unit #1451 was recorded from Monkey B, session 13.

**Fig. S9E.**
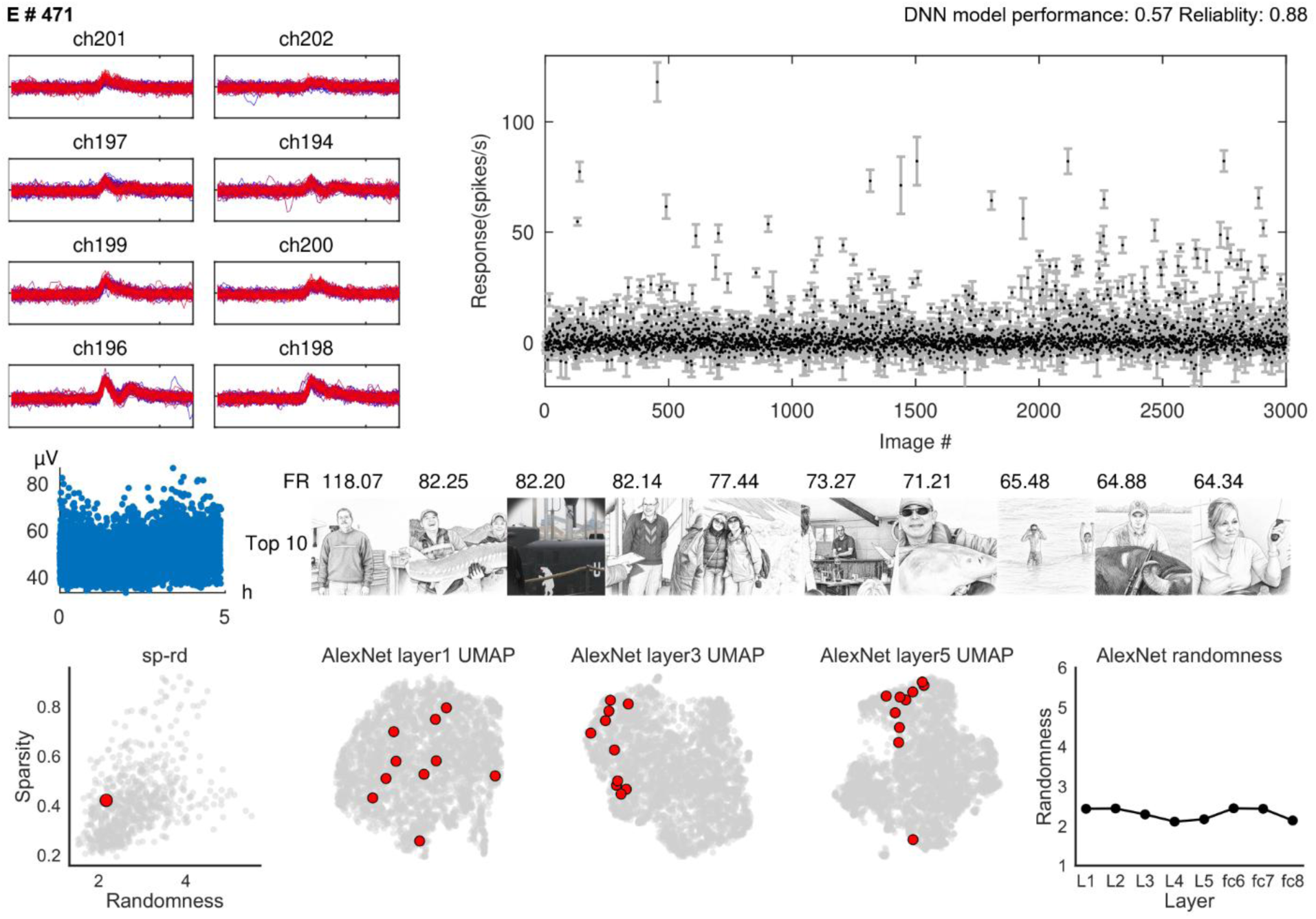
Detailed response properties of example SUA #471. Multichannel spike waveforms (upper left), mean responses across 3,000 images (upper right), spike amplitude over recording time (middle left), and the ten most effective images with their mean firing rates (middle right). Human face images have been replaced with cartoon illustrations. DNN prediction performance and response reliability are reported above the response profile. The bottom row shows the unit’s position in the joint sparseness-randomness distribution, the locations of its preferred images in UMAP embeddings derived from AlexNet layers 1, 3, and 5, and its feature-randomness values across AlexNet layers. Response points and error bars indicate the mean ± s.e.m. across repeated image presentations. Unit #471 was recorded from Monkey A, session 4.

**Fig. S9F.**
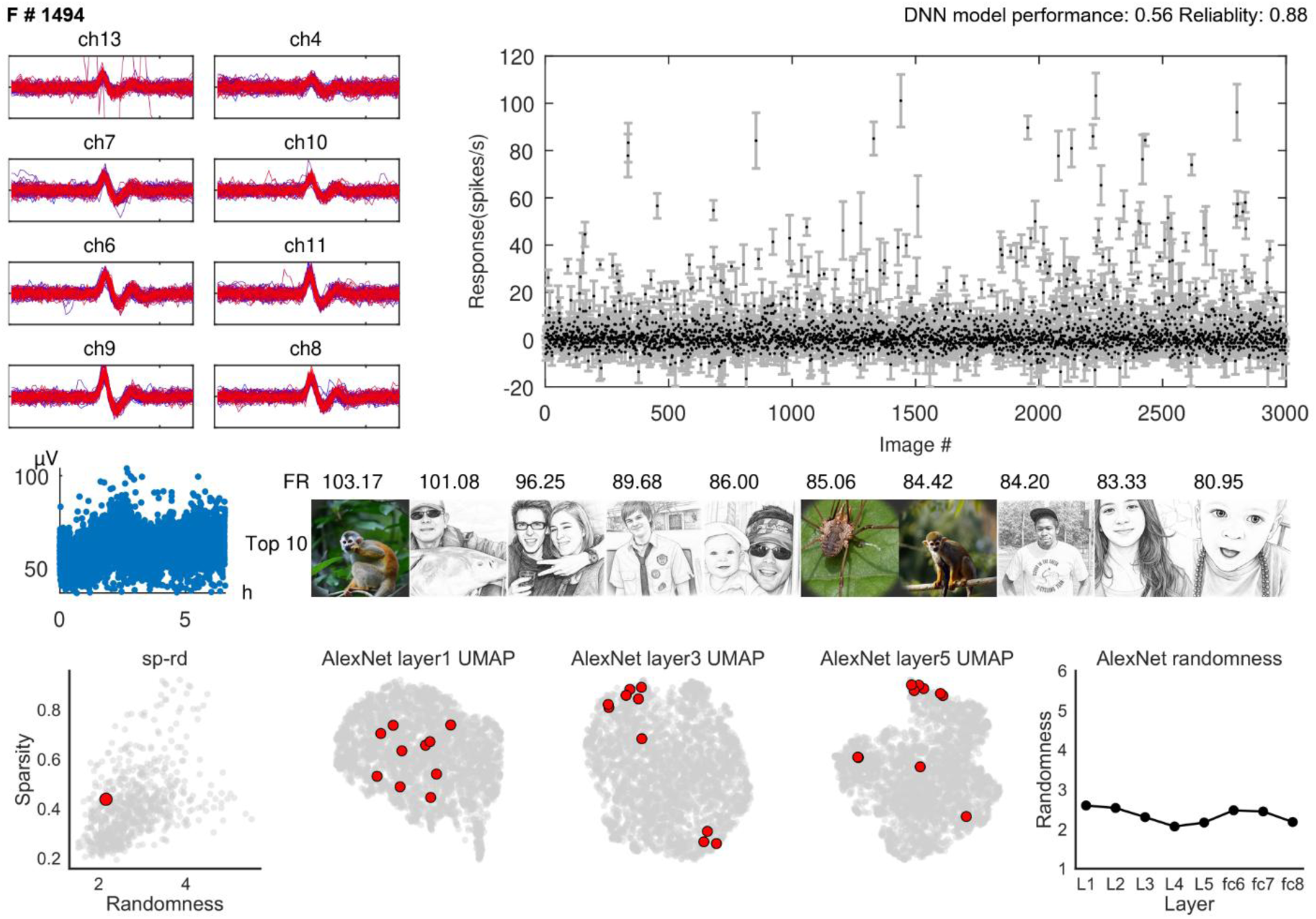
Detailed response properties of example SUA #1494. Multichannel spike waveforms (upper left), mean responses across 3,000 images (upper right), spike amplitude over recording time (middle left), and the ten most effective images with their mean firing rates (middle right). Human face images have been replaced with cartoon illustrations. DNN prediction performance and response reliability are reported above the response profile. The bottom row shows the unit’s position in the joint sparseness-randomness distribution, the locations of its preferred images in UMAP embeddings derived from AlexNet layers 1, 3, and 5, and its feature-randomness values across AlexNet layers. Response points and error bars indicate the mean ± s.e.m. across repeated image presentations. Unit #1494 was recorded from Monkey B, session 13.

**Fig. S9G.**
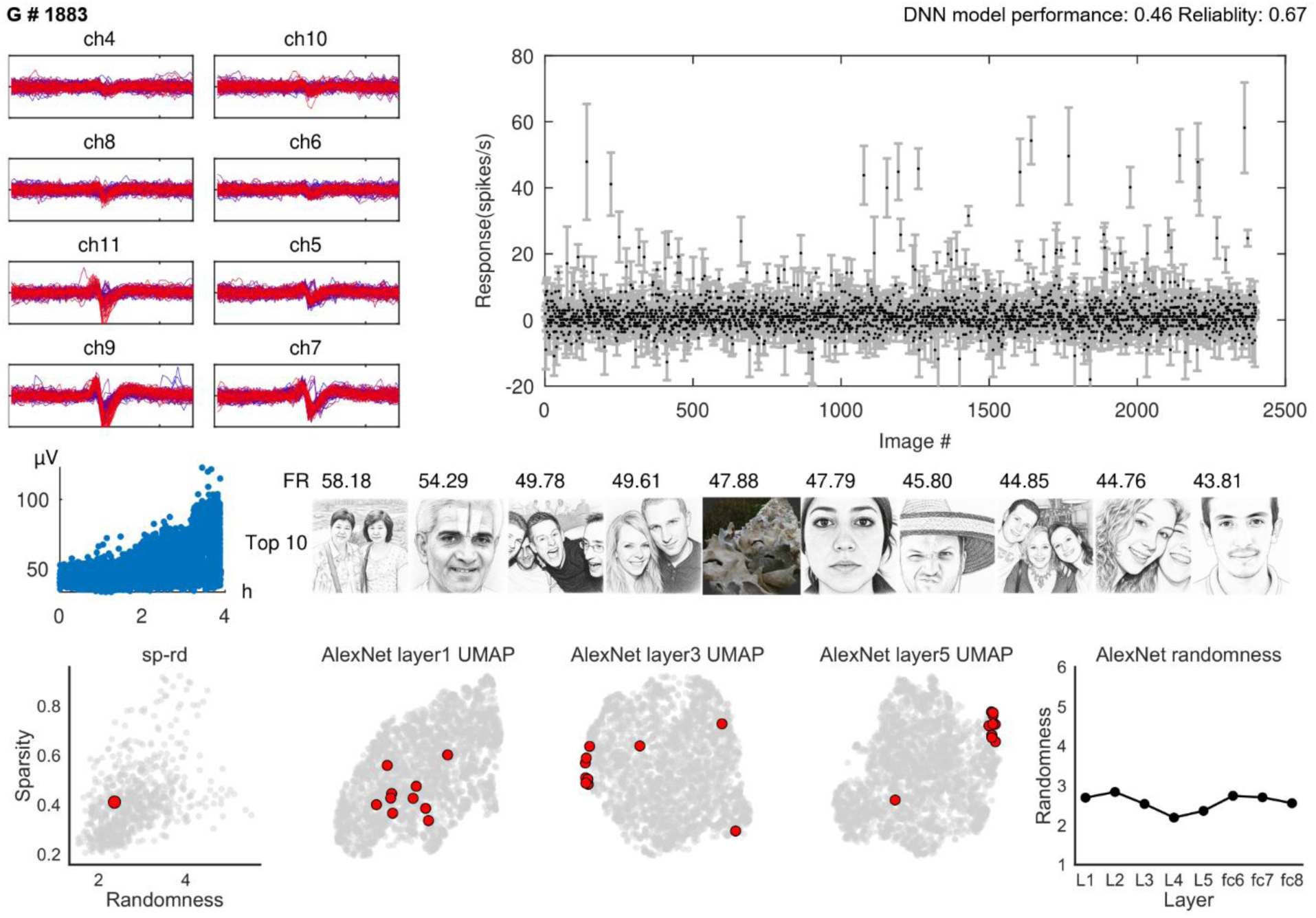
Detailed response properties of example SUA #1883. Multichannel spike waveforms (upper left), mean responses across 3,000 images (upper right), spike amplitude over recording time (middle left), and the ten most effective images with their mean firing rates (middle right). Human face images have been replaced with cartoon illustrations. DNN prediction performance and response reliability are reported above the response profile. The bottom row shows the unit’s position in the joint sparseness-randomness distribution, the locations of its preferred images in UMAP embeddings derived from AlexNet layers 1, 3, and 5, and its feature-randomness values across AlexNet layers. Response points and error bars indicate the mean ± s.e.m. across repeated image presentations. Unit #1883 was recorded from Monkey B, session 16.

**Fig. S9H.**
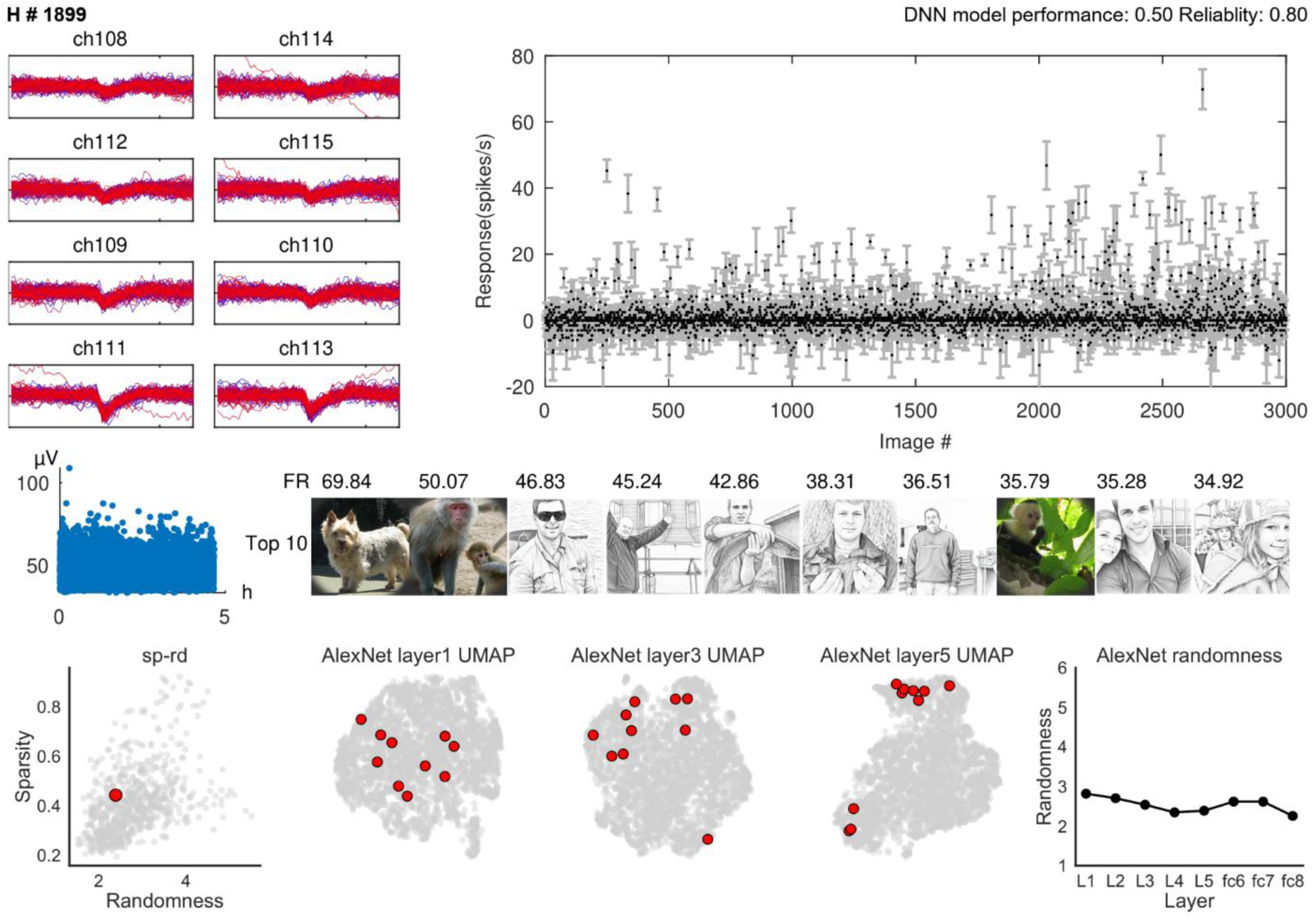
Detailed response properties of example SUA #1899. Multichannel spike waveforms (upper left), mean responses across 3,000 images (upper right), spike amplitude over recording time (middle left), and the ten most effective images with their mean firing rates (middle right). Human face images have been replaced with cartoon illustrations. DNN prediction performance and response reliability are reported above the response profile. The bottom row shows the unit’s position in the joint sparseness-randomness distribution, the locations of its preferred images in UMAP embeddings derived from AlexNet layers 1, 3, and 5, and its feature-randomness values across AlexNet layers. Response points and error bars indicate the mean ± s.e.m. across repeated image presentations. Unit #1899 was recorded from Monkey B, session 17.

**Fig. S10.**
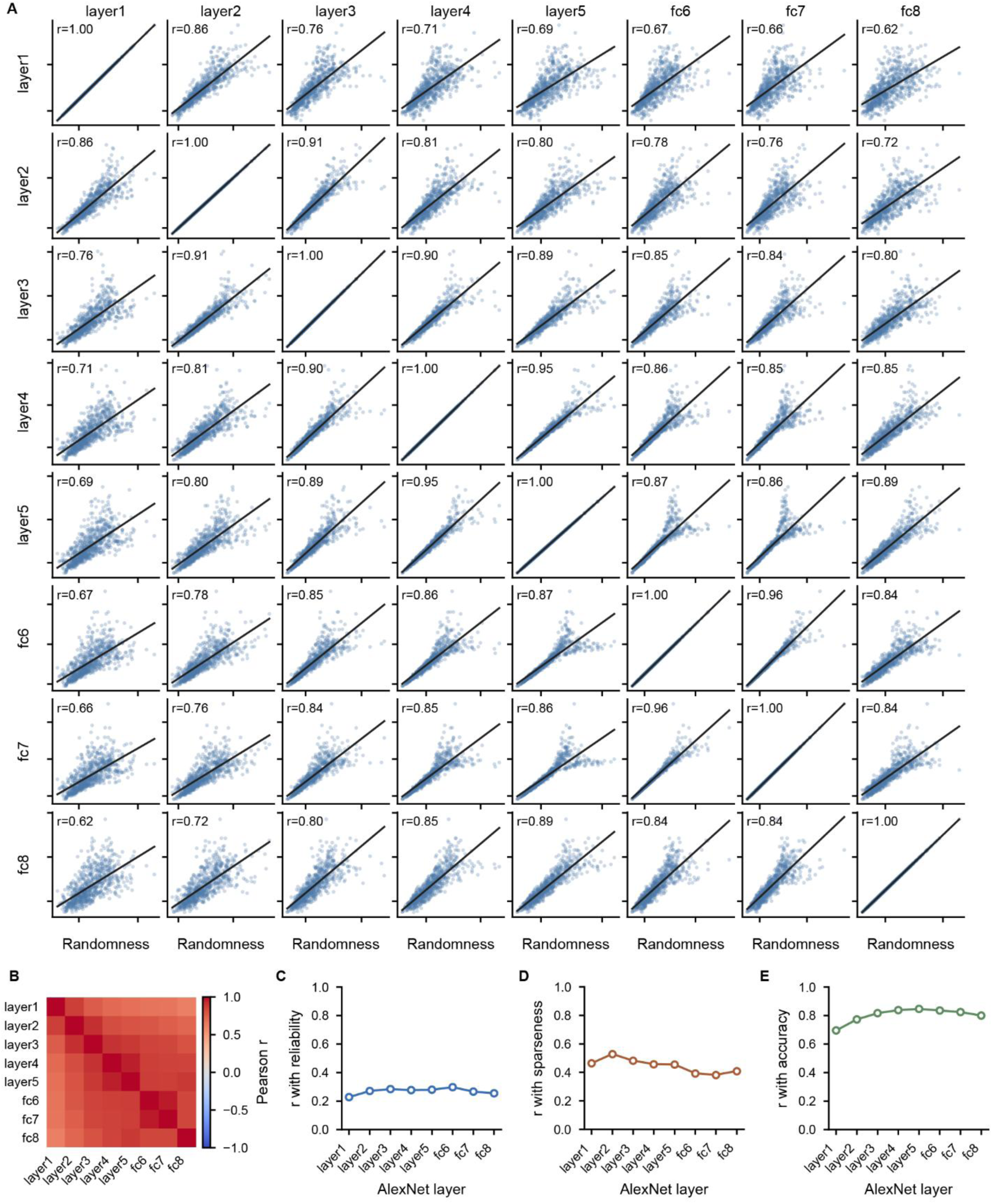
Feature-randomness estimates are consistent across AlexNet layers. **A,** Pairwise relationships between feature-randomness values computed from eight AlexNet layers across single units (*n* = 733). Each point represents one unit, black lines show linear regression fits, and reported r values are Pearson correlation coefficients. **B,** Heatmap summarizing the pairwise Pearson correlations shown in **A**. **C–E,** Pearson correlations of layer-specific feature randomness with response reliability (**C**), response sparseness (**D**), and noise-ceiling-normalized DNN prediction performance (**E**). Each point represents one AlexNet layer.

**Fig. S11.**
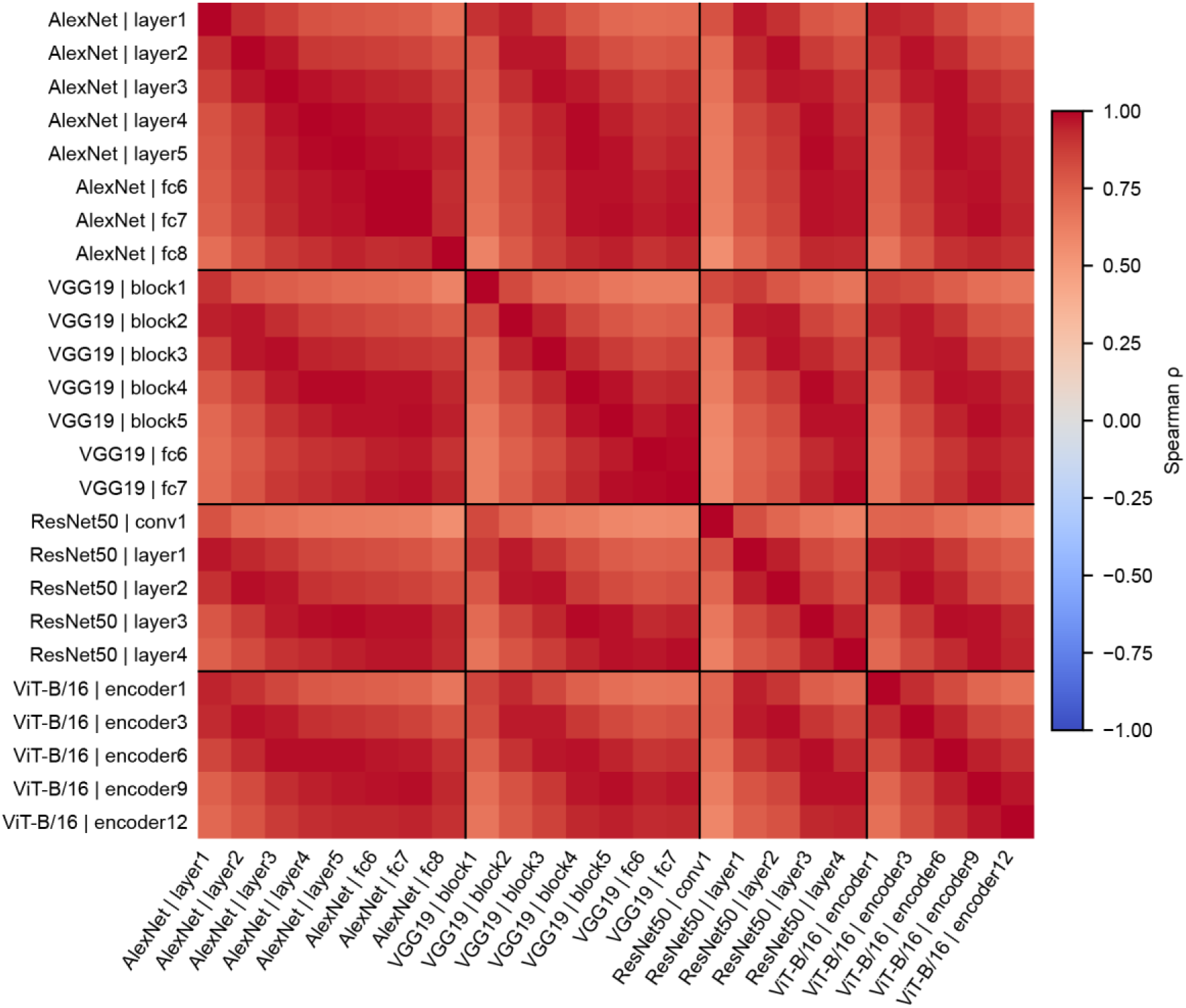
Feature-randomness estimates are consistent across DNN architectures and layers. Pairwise Spearman correlations across single units between feature-randomness estimates derived from selected layers of AlexNet, VGG19, ResNet-50, and ViT-B/16. Black lines separate the four architectures, and color indicates Spearman’s *ρ*.

**Fig. S12.**
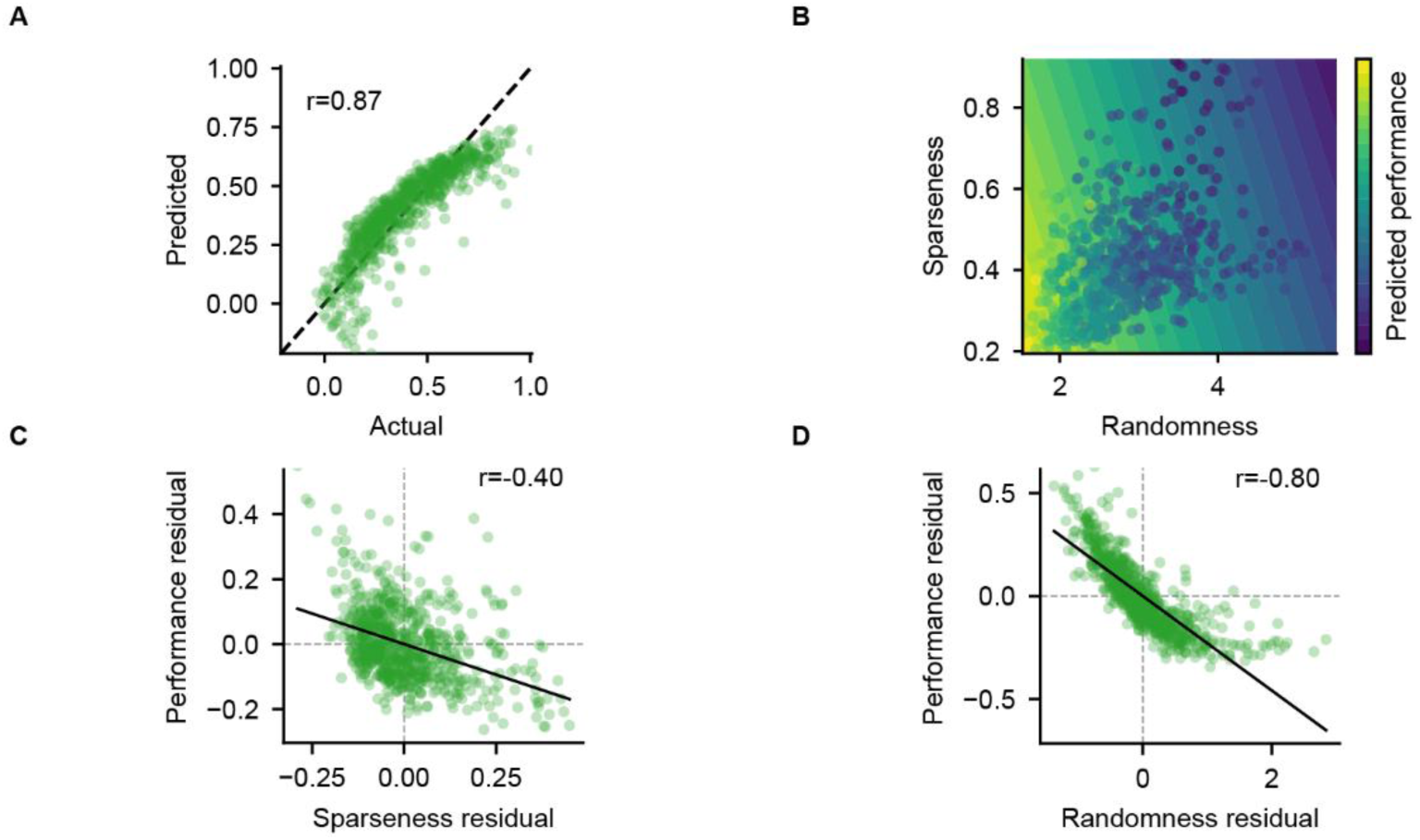
Sparseness and feature randomness make partly independent contributions to DNN prediction performance. **A,** Predicted versus observed DNN prediction performance from a two-predictor linear regression model incorporating response sparseness and feature randomness. Each point represents one unit, the dashed line indicates equality, and *r* is the Pearson correlation between predicted and observed performance. **B,** Model-predicted performance across the joint sparseness-randomness space; background contours indicate equal predicted performance. **C,** Relationship between residual sparseness and residual DNN performance after removing their linear associations with feature randomness. **D,** Relationship between residual feature randomness and residual DNN performance after removing their linear associations with sparseness. In **C** and **D**, solid lines show linear regression fits, dashed lines indicate zero, and *r* values are Pearson correlation coefficients.

**Fig. S13.**
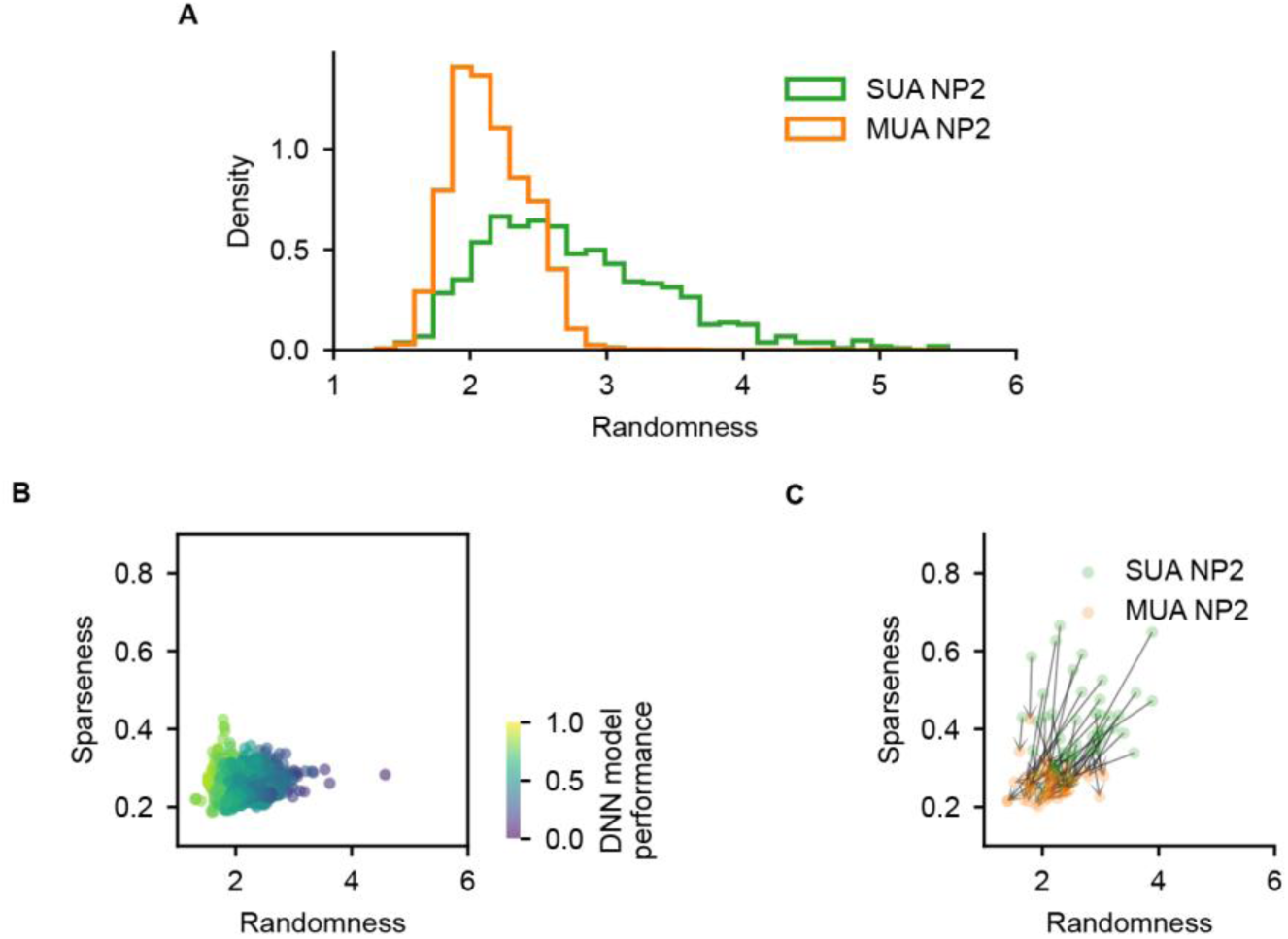
Local pooling reduces feature randomness and response sparseness. **A,** Density-normalized distributions of feature randomness for NP2 SUA and NP2 MUA. **B,** Joint distribution of response sparseness and feature randomness for NP2 MUA. Each point represents one MUA signal, and color indicates noise-ceiling-normalized DNN prediction performance. **C,** Fifty randomly sampled, spatially corresponding SUA-MUA pairs in the sparseness-randomness space. Arrows point from SUA (green) to MUA (orange).

**Fig. S14.**
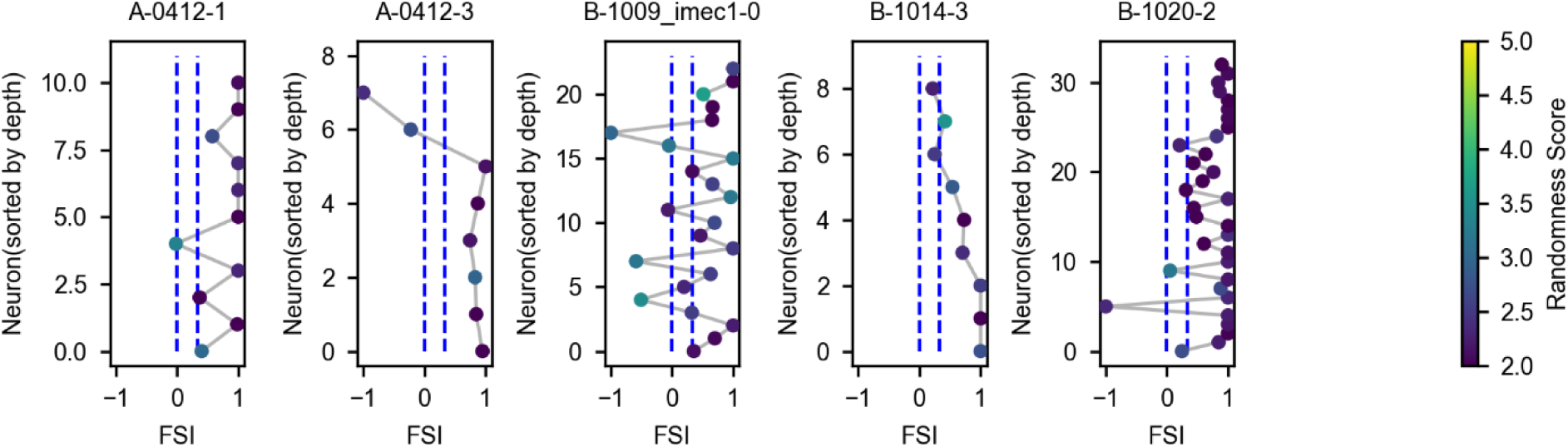
Spatial distribution of feature randomness and face selectivity. Units from five example recording sessions are ordered by recording depth along the y-axis and plotted against their face-selectivity index (FSI) along the x-axis. Gray lines connect neighboring units in depth order, and color indicates feature randomness. Blue dashed lines mark FSI = 0 and the criterion used to define face-selective units.

**Fig. S15.**
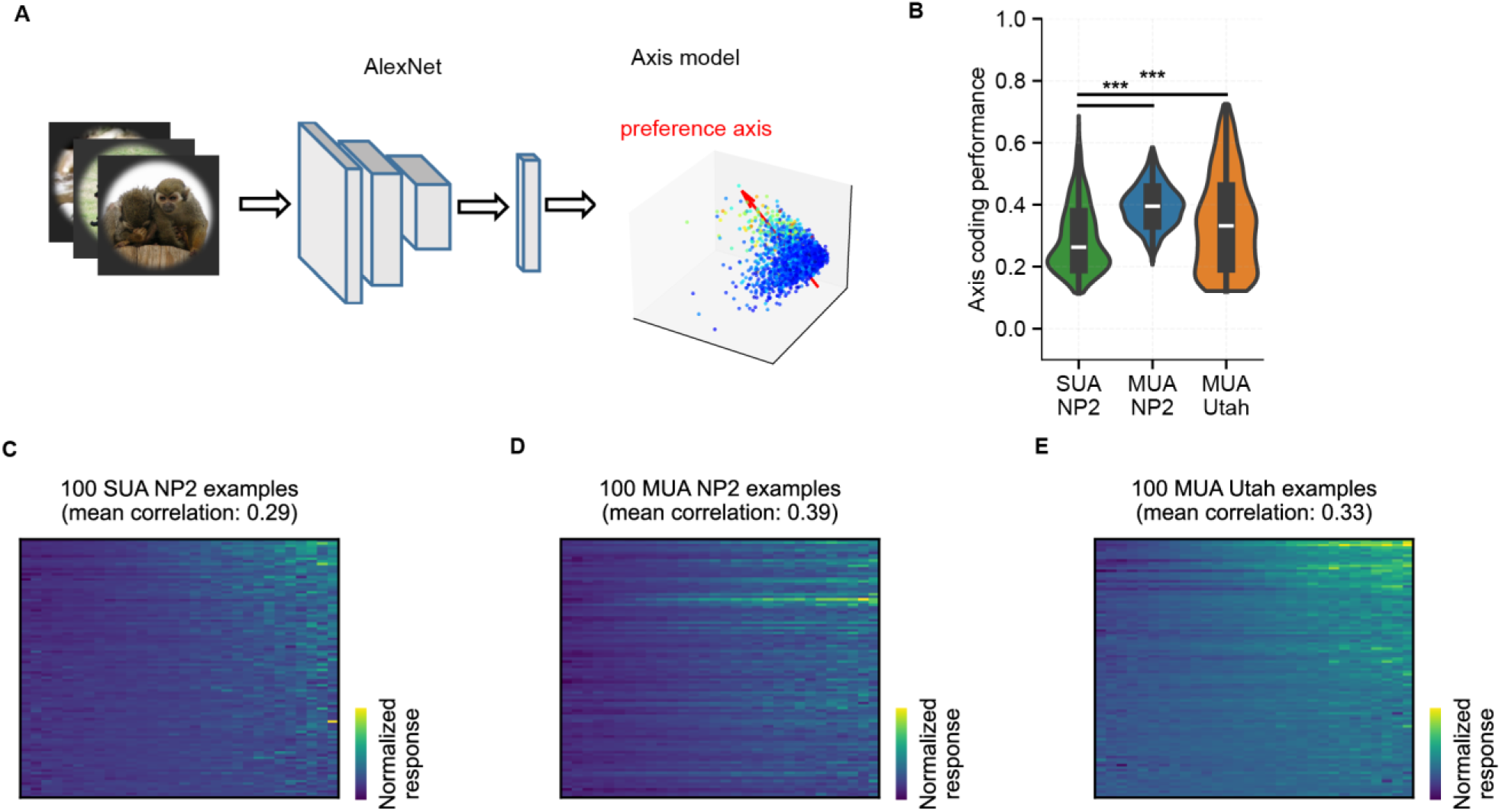
Axis-model fits to SUA and MUA responses. **A,** Schematic of the axis-coding model. Image features extracted from AlexNet were projected onto a fitted preferred axis for each response signal, and position along this axis was used to predict the signal’s image responses. **B,** Axis-model prediction performance for NP2 SUA, NP2 MUA, and Utah-array MUA. Black boxes indicate the median and interquartile range. Horizontal bars indicate significant comparisons (\*\*\**P* < 0.001; two-sided Mann–Whitney *U*-tests). **C– E,** Normalized responses of 100 randomly sampled NP2 SUA (**C**), NP2 MUA (**D**), and Utah-array MUA signals (**E**), with images ordered for each signal according to their projection onto its fitted preferred axis. Mean response-axis correlations are reported above each matrix.

**Fig. S16.**
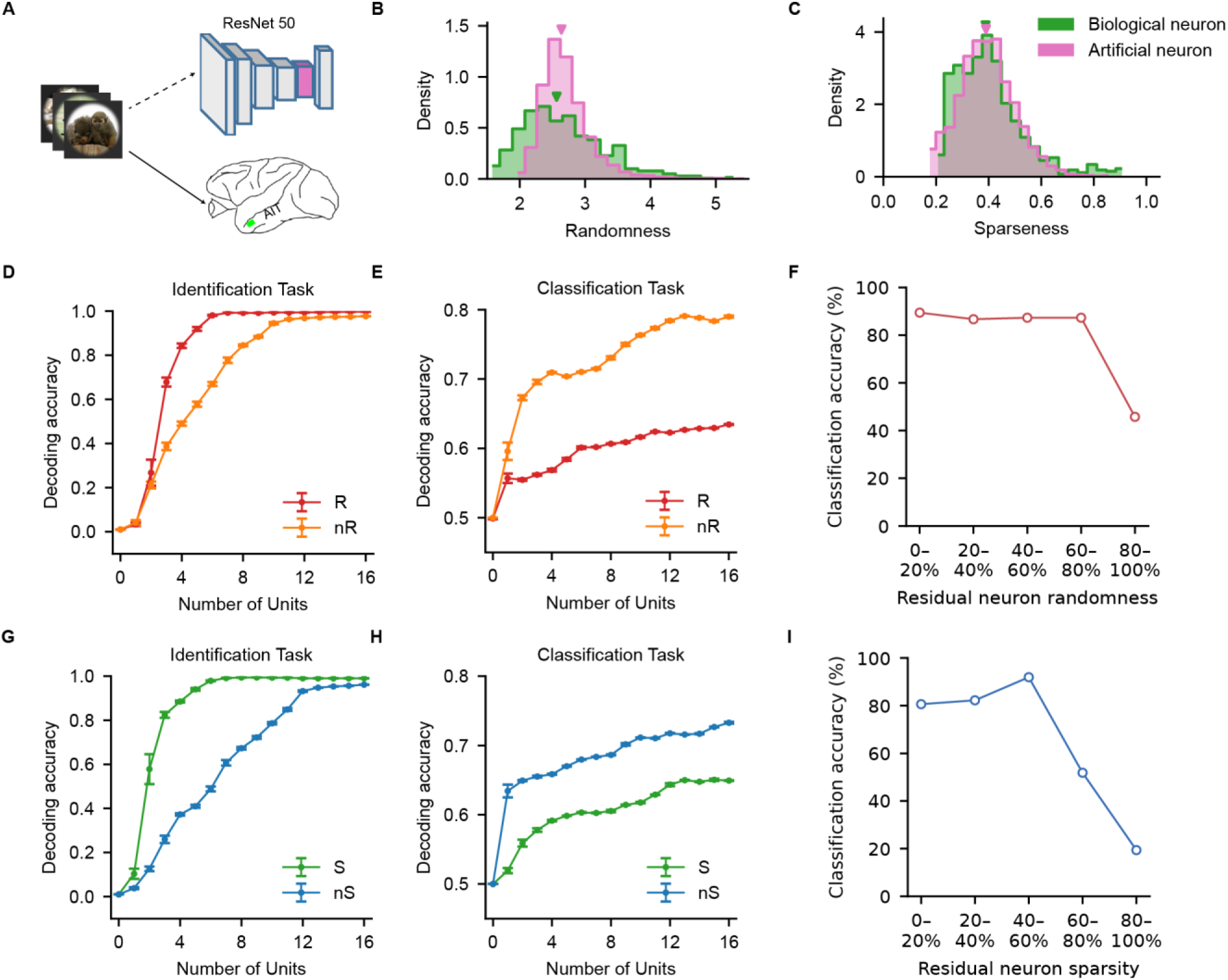
Feature randomness and response sparseness organize identification and classification in ResNet-50 units. **A,** Comparison of artificial ResNet-50 units and biological AIT single units. Five thousand artificial units were randomly sampled from 154 ResNet-50 layers after excluding units whose maximum activation across the experimental images did not exceed 10. **B, C,** Density distributions of feature randomness (**B**) and response sparseness (**C**) for artificial and biological units. Arrowheads indicate medians. **D, E,** Image-identification (**D**) and face-versus-non-face classification (**E**) performance as a function of the number of high-randomness (R) or low-randomness (nR) artificial units included in the decoder. **F,** Network classification accuracy after retaining artificial units from successive quintiles of feature-randomness residuals after controlling for sparseness. **G, H,** Image-identification (**G**) and face-versus-non-face classification (**H**) performance as a function of the number of sparse (S) or non-sparse (nS) artificial units included in the decoder. Curves and error bars in **D**, **E**, **G**, and **H** indicate the mean ± s.e.m. across repeated random subsamples. **I,** Network classification accuracy after retaining artificial units from successive quintiles of sparseness residuals after controlling for feature randomness.

**Fig. S17.**
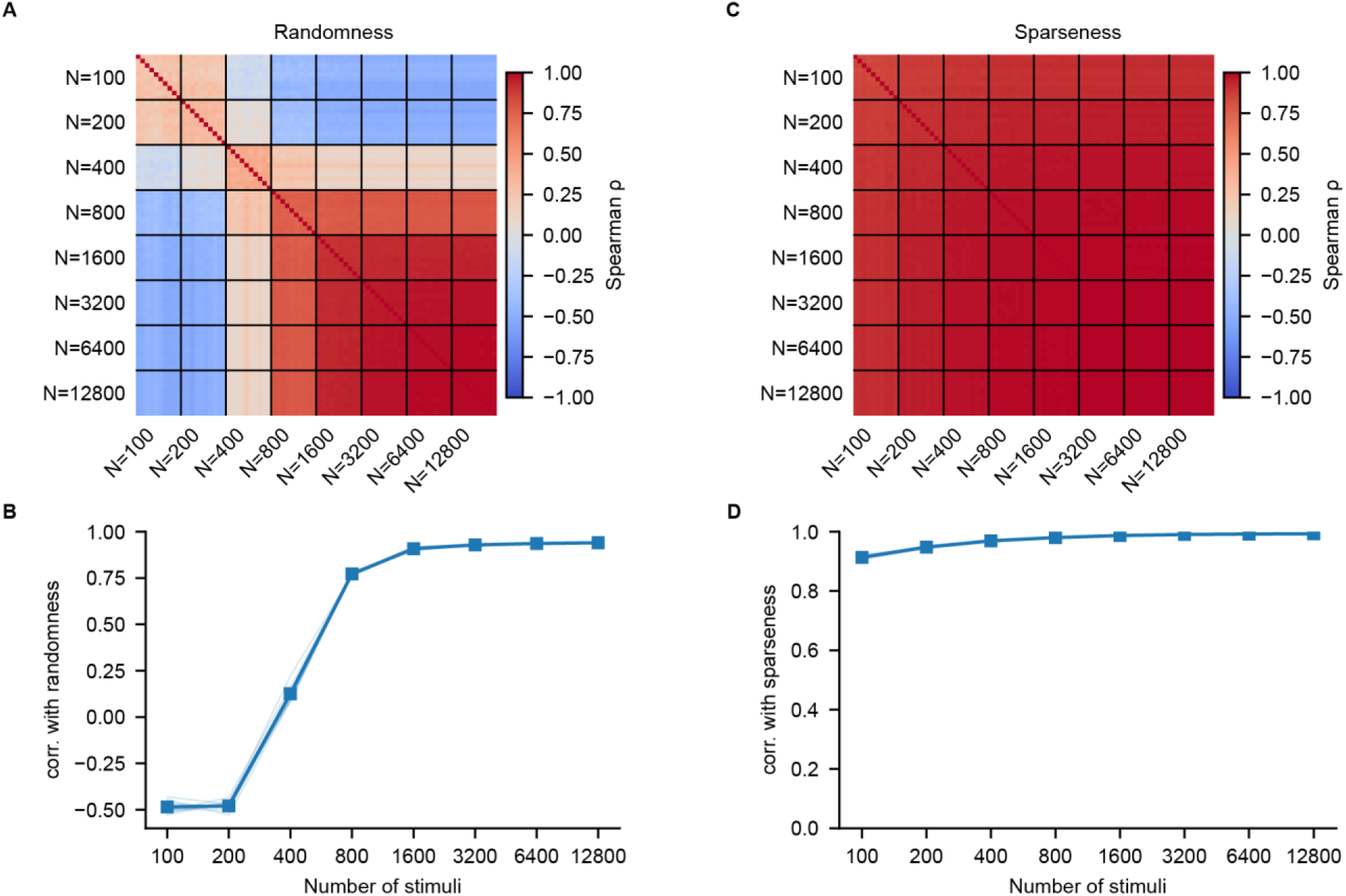
Sensitivity of feature-randomness and sparseness estimates to stimulus sample size. Feature randomness and response sparseness were estimated for artificial ResNet-50 units using randomly sampled image sets ranging from 100 to 12,800 images. Sampling was repeated ten times at each sample size. **A,** Pairwise Spearman correlations between feature-randomness estimates obtained from all sampled image sets. Black lines delineate sample sizes, and each row and column represents one random sample. **B,** Correlation between feature-randomness estimates obtained at each sample size and estimates obtained using the experimental image set. **C,** Pairwise Spearman correlations between response-sparseness estimates obtained from all sampled image sets. **D,** Correlation between response-sparseness estimates obtained at each sample size and estimates obtained using the experimental image set. In **B** and **D**, thin lines show individual sampling repetitions and the blue line with square markers shows the mean.

